# The CoREST Repressor Complex Mediates Phenotype Switching and Therapy Resistance in Melanoma

**DOI:** 10.1101/2020.09.30.320580

**Authors:** Muzhou Wu, Ailish Hanly, Frederick Gibson, Kevin Kuang, Jay Kalin, Sarah Nocco, Marianne Collard, Matthew Cole, Amy Xiao, Filisia Agus, Adam Labadorf, Philip A Cole, Rhoda M. Alani

## Abstract

Virtually all patients with BRAF-mutant melanoma develop resistance to MAPK inhibitors largely through non-mutational events^1,2^. Although the epigenetic landscape has been shown to be altered in therapy-resistant melanomas and other cancers^3,4^, a specific targetable epigenetic mechanism regulating treatment resistance has not been validated to date. Here we evaluate the CoREST repressor complex and the novel inhibitor, corin^5^, within the context of melanoma phenotype plasticity and therapeutic resistance in order to define epigenetic mechanisms underlying these processes. We find that CoREST is a critical mediator of the major distinct melanoma phenotypes and that corin treatment of melanoma cells leads to phenotype reprogramming. We further demonstrate that treatment of BRAF inhibitor (BRAFi)-resistant melanomas with corin leads to resensitization of tumor cells to BRAFi. Among the transcriptional targets of CoREST in melanoma are the dual-specificity phosphatases (DUSPs). DUSP1 is shown to be consistently downregulated in BRAFi-resistant melanomas which can be reversed by corin treatment, thereby leading to downstream inhibition of p38 MAPK activity and resensitization of resistant cells to targeted BRAFi therapies. These findings identify the CoREST repressor complex as a central mediator of melanoma phenotype plasticity and resistance to targeted therapy and suggest that CoREST inhibitors may prove beneficial to patients with BRAF-mutant melanomas who have acquired BRAFi-resistance.

## Main

Melanomas exhibit phenotype plasticity which allows them to switch between distinctive transcriptional programs in response to external stressors, including targeted therapies^4^. These transcriptional phenotypes are characterized by altered differentiation and metabolic states including a proliferative/differentiated/MITF^high^/AXL^low^ phenotype and an undifferentiated/invasive/MITF^low^/AXL^high^phenotype^6,7^ with associated changes in the epigenetic landscape^8–9^. Moreover, cellular plasticity is a driver of resistance to targeted therapies in melanoma and other cancers, with dynamic transitions between distinctive molecular phenotypes promoting MAPKi bypass mechanisms^10^. As reversible transcriptional reprogramming dictates the plasticity of molecular phenotypes, research has focused on the role of epigenetic regulation in this process^11–13^. The CoREST repressor complex is a member of the class 1 histone deacetylase family of repressor complexes that was originally identified as a cofactor for REST repression and regulation of neuron-specific gene silencing during development^14^. RCOR1 functions as a scaffold for the CoREST repressor complex promoting crosstalk between HDAC1/2 and the H3K4me demethylase LSD1^15^, and has recently been shown to regulate Treg function and antitumor immunity^16^. We recently described a potent and specific dual-warhead inhibitor of the CoREST complex targeting HDAC1/2 and LSD1, corin, that demonstrates growth inhibition in melanoma, cutaneous squamous cell carcinoma^5^, and diffuse intrinsic pontine glioma.^17^ We therefore hypothesized that CoREST inhibition may elicit synergistic growth inhibition with BRAFi therapies through epigenetic reprogramming of molecular phenotypes in BRAF-mutant melanomas.

### CoREST inhibition mediates phenotype switching in melanoma and resensitizes BRAF-inhibitor resistant melanoma cells to BRAF inhibitor therapy

To further explore the role of CoREST in human melanoma development, we treated a panel of phenotypically distinct melanoma cell lines with corin for 24 h (Fig. 1a-e). Corin treatment of MITF^high^/AXL^low^ melanoma cells led to decreased expression of MITF and increased expression of histone H3K4me2 and H3K9ac/K27ac marks without significant effects on the MEK/ERK pathway (Fig. 1a). Additionally, corin treatment of MITF^low^/AXL^high^ melanoma cells led to decreased AXL expression and increased histone acetylation and methylation marks without impacting the MEK/ERK pathway (Fig. 1b), suggesting a specific reversion of melanoma differentiation phenotypes to intermediate cellular programs. Corin treatment inhibited growth in the majority of MITF^high^/AXL^low^ melanoma cells; however, minimal growth inhibition and even enhanced cellular proliferation was seen in MITF^low^/AXL^high^ cells treated with corin for 24 h (Fig. 1c), consistent with conversion to an intermediate proliferation phenotype following corin treatment. Morphological changes were also observed following corin treatment, with MITF^high^/AXL^low^ cells demonstrating a more epithelioid cellular phenotype, while MITF^low^/AXL^high^ cells demonstrated a more elongated/differentiated phenotype (Extended Data Fig. 1a, b). In addition, corin treatment increased cellular invasion and expression of focal adhesions in MITF^high^/AXL^low^ cells while decreasing invasion and expression of focal adhesions in MITF^low^/AXL^high^ cells (Figs. 1d, e). As phenotype switching is associated with resistance to targeted therapies in melanoma and other cancers and intermediate phenotypes are generally treatment-sensitive, we next investigated the effect of CoREST inhibition combined with the BRAFi, PLX4032 (vemurafenib) on BRAFi-resistant (BRAFi-R) melanoma cell proliferation. Notably, we found that corin significantly increased antiproliferative effects of PLX4032 in BRAFi-R melanoma cells (Fig. 1f). In addition, PLX4032 enhanced the antiproliferative effects of corin alone, with strong synergy identified between PLX4032 and corin (combination index <0.1^18^) versus the LSD1 inhibitor (compound 7) or HDAC1 inhibitor (MS275) alone (Extended Data Fig. 1c, d, e). Corin treatment also significantly reduced colony formation and increased apoptosis in BRAFi-R melanoma cells, which was further enhanced in combination with PLX4032 (Figs. 1g,1h).

**Figure 1.**
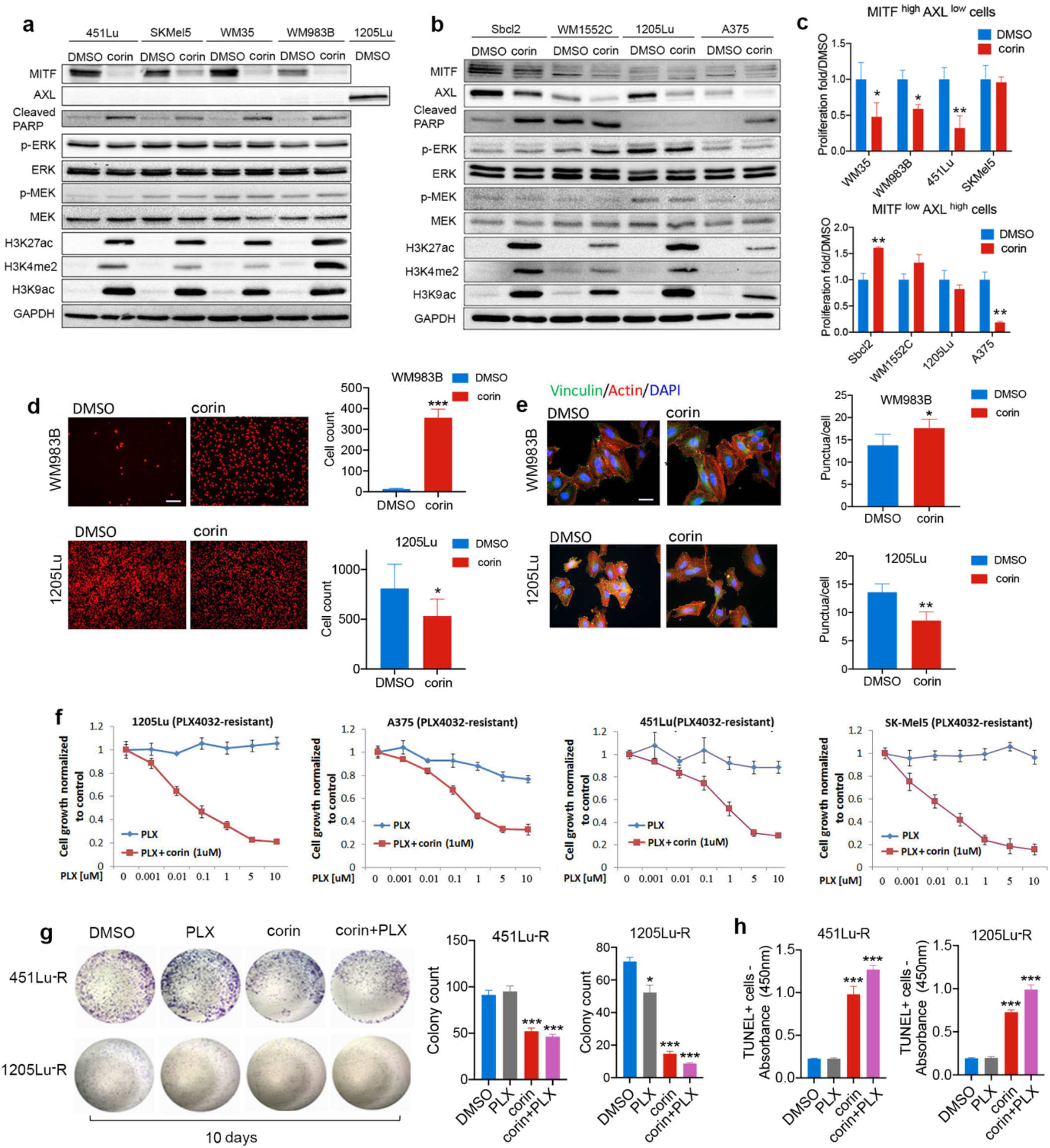
Corin mediates phenotype switching in melanoma cell lines and promotes resensitization of BRAFi-resistant cells to BRAF inhibitor treatment. **a, b,** Western blot analysis in MITF^high^/AXL^low^ cell lines 451Lu, SKMel5, WM35 and WM983B **(a)** and MITF^low^/AXL^high^ cell lines Sbcl2, WM1552C, 1205Lu and A375 **(b)** following 24 h treatment with DMSO or 2.5 μM corin. **c**, Cellular proliferation of MITF^high^/AXL^low^ (upper panel) and MITF^low^/AXL^high^ (lower panel) cell lines following 24 h treatment with DMSO or 2.5 μM corin. **d,** Invasion assay and quantification MITF^high^/AXL^low^ melanoma cells (WM983B, upper panel) and MITF^low^/AXL^high^ melanoma cells (1205Lu, lower panel) following 24 h treatment with DMSO or 2.5 μM corin (representative images shown, scale bar = 100 μm). **e,** Vinculin staining of focal adhesions in WM983B (upper panel) and 1205Lu (lower panel) following 24 h treatment with DMSO or 2.5 μM corin (representative images shown, scale bar = 20 μm). **f,** Proliferation assays of 1205Lu and A375 (MITF^low^/AXL^high^) BRAFi-resistant cell lines, and SK-Mel5, 451Lu (MITF^high^/AXL^low^) BRAFi-resistant cell lines treated with increasing doses of PLX4032 +/− 1μM corin for 72 h. **g**, Colony formation assay and quantification of colony counts in 451Lu-R and 1205Lu-R cells treated with DMSO, PLX4032 alone, corin alone, and corin + PLX4032. **h,** Quantification of TUNEL-positive cells in 451Lu-R and 1205Lu-R cells following 72 h treatment with DMSO, PLX4032 alone, corin alone, and corin + PLX4032. Data (mean ± SE) are representative of at least two independent experiments (***p < 0.001, **p <0.01, *p <0.05).

### CoREST complex inhibition promotes expression of DUSPs and inhibits p38 MAPK in BRAFi-resistant melanoma cells

To investigate mechanisms of corin resensitization of BRAFi-R melanoma cells to PLX4032, genome-wide RNA sequencing was performed on 451Lu-R (MITF^high^/AXL^low^) and 1205Lu-R (MITF^low^/AXL^high^) cells treated with corin for 24 h. Differentially regulated genes were identified by comparison with DMSO controls (Fig. 2a). Not surprisingly, corin treatment led to significantly greater numbers of up-regulated genes than down-regulated, consistent with the repressive function of the CoREST complex. Both up- and down-regulated gene sets between the two cell lines possess significant numbers of common corin-affected genes as well as cell line-specific gene sets (Fig 2a, Extended Data Fig. 2a). Differentially regulated genes identified by RNA-Seq were further subjected to Ingenuity Pathway Analysis (IPA) and analyzed by gene set enrichment analysis (GSEA) using the Kyoto Encyclopedia of Genes and Genomes (KEGG) pathway gene sets. Corin-induced common enriched pathways in 451Lu-R and 1205Lu-R cells included MAPK signaling, Hippo signaling, cell adhesion molecules, axon guidance, Wnt signaling, cell cycle regulation, interferon gamma and PD-1 signaling as well as distinct pathways specific to each cell line (Figs. 2b, c, Extended Data Fig. 2b-i, Extended Data Fig. 3). Comparison of corin-associated RNA-Seq data with publicly available datasets^6,19,20^ supported a phenotype switch signature following corin treatment in both 451Lu-R and 1205Lu-R cells (Fig. 2d). Genes associated with the corin-induced phenotype switch included AXL, MITF, SOX10, WNT5A, PAX3, ZEB1, ZEB2, PGC1a, DUSP1, and DUSP5 (Fig. 2d). Interestingly, the dual-specificity phosphatases, DUSP1 and DUSP5, were among the genes whose expression was most notably altered in the MAPK signaling GSEA and significantly upregulated following corin treatment in both BRAFi-R cell lines (Fig. 2d, e). In addition, corin treatment increased expression of type 1 and type 2 interferon response genes and repetitive elements (Extended Data Fig. 4a-c), and decreased expression of RNA-induced silencing complex (RISC) components (Extended Data Fig. 4d), similar to previously reported changes associated with LSD1 ablation in tumor cells^21^.

**Figure 2.**
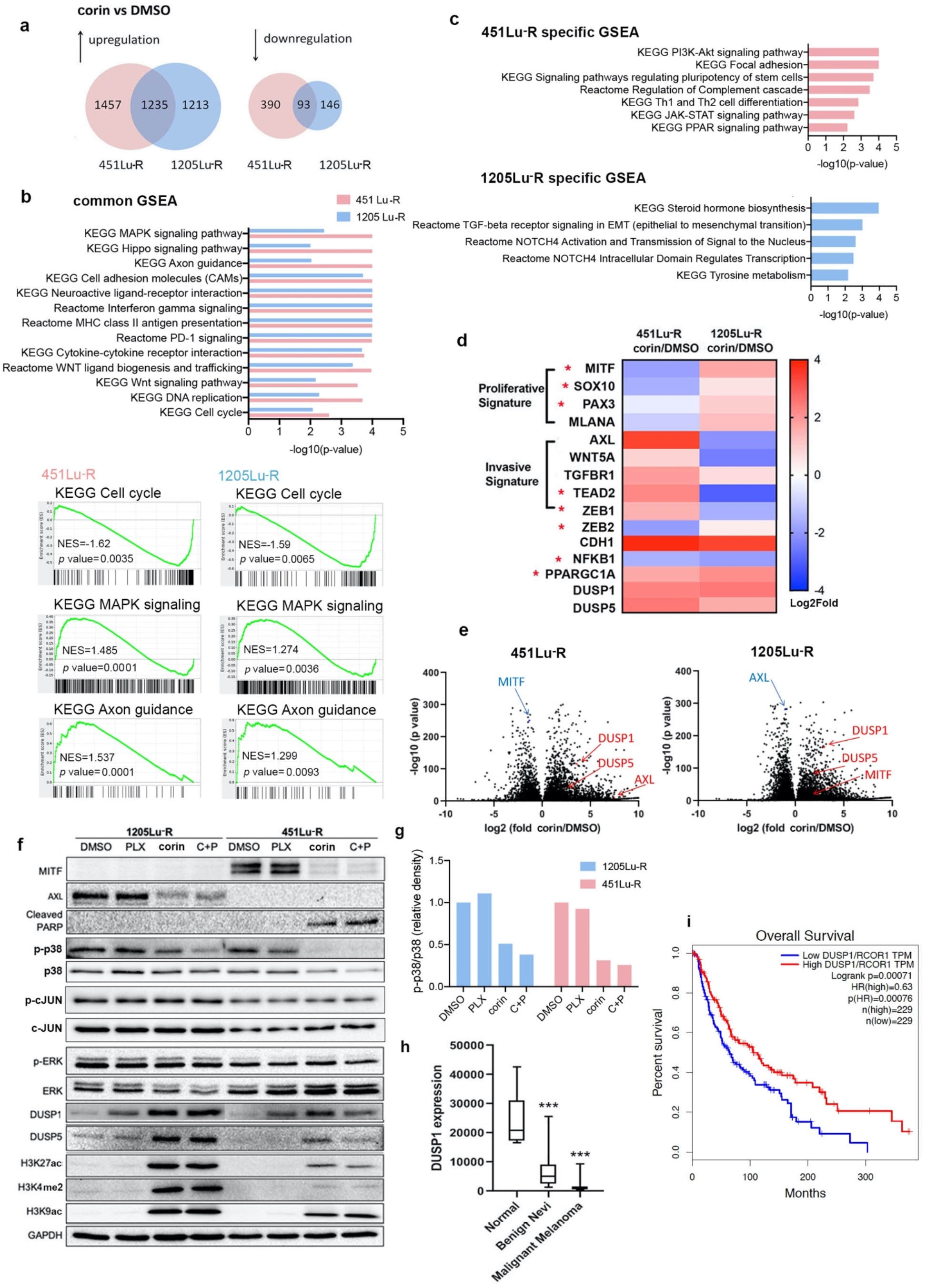
Corin promotes phenotype switching and melanoma re-sensitization to BRAFi therapy through DUSP1-induced inhibition of p38 MAPkinase. **a,** RNA sequencing profiling of 451Lu-R and 1205Lu-R following 24 h treatment with DMSO versus 2.5 μM corin. Venn Diagram illustrates corin-induced upregulated and downregulated expression of cell line-specific and common genes (fold change ≥ 4σ, FDR < 0.001). **b,** IPA analysis of corin-induced common enriched pathways in 451Lu-R and 1205Lu-R. GSEA in corin-versus DMSO-treated samples illustrating representative common gene sets in 451Lu-R and 1205Lu-R. NES (Normalized Enrichment Score), (FDR < 0.05). **c,** IPA analysis of corin-induced distinct pathways in 451Lu-R and 1205Lu-R. **d,** Heatmap of differential expression patterns of proliferative vs. invasive gene signatures, as well as transcriptional regulators associated with distinct melanoma phenotypes (depicted by * in red). The Log2 fold change difference is color-coded (red, upregulated; blue, downregulated). **e,** Volcano plot of transcript changes in 451Lu-R and 1205Lu-R following corin treatment with notable changes in DUSP1, DUSP5, MITF and AXL. **f,** Western blot analysis of 1205Lu-R and 451Lu-R cells treated with DMSO, PLX4032 alone, corin alone, or corin + PLX4032 for 24 h. **g,** Quantification of relative expression of pp38 (active) versus p38 (total) protein expression in 1205Lu-R and 451Lu-R melanoma cells following treatment with DMSO, PLX4032 alone, corin alone, or corin + PLX4032. **h**, Quantification of DUSP1 expression in normal skin, benign nevi, and malignant melanoma patient tissues. **i,** Kaplan-Meier curves illustrating the relationship between the expression of DUSP1/RCOR1 in patient tumor specimens and overall patient survival. All data (mean ± SE) are representative of at least two independent experiments (***p <0.001).

To further investigate mechanisms of corin resensitization of BRAFi-R melanoma cells to PLX4032, RNA sequencing was performed on 451Lu-R and 1205Lu-R cells treated with PLX4032 +/− corin. Differentially regulated genes were identified by comparison with PLX4032-treated cells and further subjected to IPA analysis. Similarly enriched pathways and GO terms were identified in these transcriptomic comparisons as in corin vs. DMSO controls (Extended Data Fig. 5, Extended Data Fig. 6). MAPK signaling pathways, cell adhesion and Wnt signaling pathways, as well as pathways associated with DNA replication and DNA repair were most significantly perturbed upon corin + PLX4032 treatment in both cell phenotypes (Extended Data Fig. 5c). Interestingly, genes previously noted to be downregulated in patient tumors following acquired MAPKi-resistance^22^ were found to be specifically upregulated following corin + PLX4032 treatment of 451Lu-R and 1205Lu-R cells (Extended Data Figure 7).

Western blot analysis of BRAFi-R cell lines treated with corin showed reduced MITF expression in MITF^high^/AXL^low^ cells (451Lu-R) and reduced AXL expression in MITF^low^/AXL^high^ cells (1205Lu-R), consistent with a corin-associated phenotype switch (Fig. 2f, Extended Data Fig. 8a, b) which is retained in the setting of PLX4032 treatment. Corin also induced expression of H3K4me2, H3K9ac, and H3K27ac histone marks in BRAFi-R melanomas that were maintained following PLX treatment (Fig. 2f, Extended Data Fig. 8a, b). Additionally, corin significantly increased expression of the dual-specificity phosphatases, DUSP1 and DUSP5, in both BRAFi-R cell lines (Fig. 2f), consistent with the RNA-seq findings (Fig. 2d,e). As DUSPs are phosphatases that negatively regulate MAPK activity, we sought to further explore the functional significance of DUSP1 and DUSP5 as downstream effectors of corin, with regard to MAPK family members ERK, JNK and p38 MAPK, which have all been implicated in melanoma resistance to targeted therapies^23–26^. Further evaluation of the ERK MAPK signaling cascade revealed no significant effects of corin on ERK activation (Fig. 2f), suggesting that ERK-specific DUSP5 is not a critical mediator of corin-induced BRAFi resensitization in BRAFi-R cells. However, examination of corin effects on p38 MAPK and JNK/c-JUN demonstrated significantly decreased phospho-p38 levels in BRAFi-R cells, while JNK/c-Jun activity was not impacted (Fig. 2f, g, Extended Data Fig 8c, d) indicating selective DUSP1-mediated inhibition of p38 MAPK.

To further explore the relationship between DUSP1 expression and melanomagenesis, we mined publicly available gene expression datasets for normal skin, benign nevi, and primary melanoma patient tissues from The Cancer Genome Atlas (TCGA) repository. Notably, DUSP1 expression was significantly decreased in malignant melanoma compared to both benign nevi and normal skin (Fig. 2h) and decreased DUSP1 protein was noted in malignant melanoma tissues versus benign nevi by immunohistochemistry (Extended Data Fig. 9a). TCGA data was further analyzed to explore the relationship between DUSP1 and RCOR1 expression in 412 patient melanoma specimens. Kaplan-Meier plots of overall survival revealed that patients whose tumors showed higher ratios of DUSP1/RCOR1 expression had significantly increased overall survival compared to patients with lower DUSP1/RCOR1 ratios (Fig. 2i), suggesting that high DUSP1 expression in the setting of low RCOR1 expression may be predictive of improved survival in patients.

### DUSP1 is a direct target of the CoREST complex which promotes response to BRAFi therapy

To further explore the role of the CoREST complex in BRAFi-resistant melanoma, shRNA was used to stably knockdown CoREST1 in human melanoma cells (Fig 3a), which were evaluated for tumor cell growth in the presence of the BRAFi, PLX4032. CoREST1 knockdown in 451Lu-R and 1205Lu-R cells restored responsiveness to PLX4032 in a dose-dependent manner, with 50% growth reduction at the highest doses of PLX4032 compared to controls (Fig. 3b). As increased DUSP1 and DUSP5 expression occurred with corin treatment of BRAFi-R melanoma cells, we sought to establish baseline levels of DUSPs in BRAF-sensitive and BRAF-resistant melanoma cell lines. Remarkably, we found significantly reduced expression of several DUSPs, including DUSP1, in 451Lu-R and 1205Lu-R melanoma cell lines compared to their BRAFi-sensitive counterparts (Fig. 3c), suggesting a potential role of DUSP proteins in regulating BRAFi resistance. Moreover, treatment of 1205Lu-R with corin alone or combination treatment with corin and PLX4032 resulted in five-fold and eight-fold increases in DUSP1 expression, respectively, compared to vehicle (Fig. 3d), which was similarly observed in other melanoma cell lines (Extended Data Fig. 9b-d). In addition, a timecourse evaluation of DUSP1 and DUSP5 expression in four BRAFi-R melanoma cell lines treated with corin demonstrated significant upregulated expression of DUSP1 and DUSP5 within 4 hours of corin treatment (Extended Data Fig. 9e-g).

**Figure 3.**
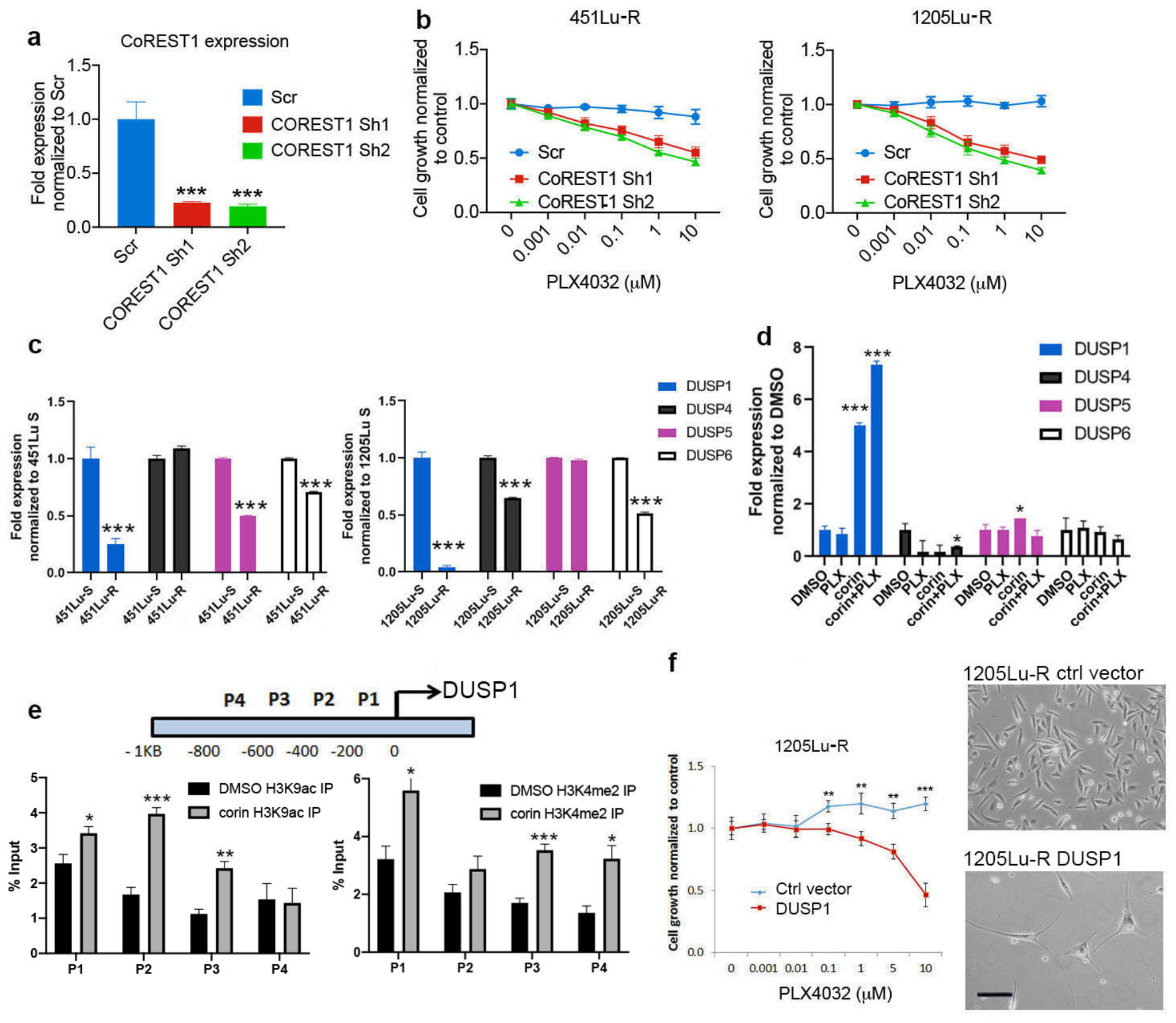
DUSP1 is a direct target of the CoREST complex and modulates p38 activity. **a,** CoREST1 knockdown by shRNA in 1205Lu-R cells, confirmed by qRT-PCR. **b**, PLX4032 dose response curves (72 h treatment) in 451Lu-R and 1205Lu-R cells following knockdown of CoREST1. **c,** DUSP1, 4, 5, and 6 expression levels (qPCR) in BRAFi-sensitive and resistant 451Lu (left) and 1205Lu (right) melanoma cells. **d,** DUSP1, 4, 5, and 6 expression in response to 24 h treatment of DMSO, PLX4032, corin, and corin + PLX4032. **e,** ChIP-PCR analyses of the occupancy of H3K9ac and H3K4me2 of the DUSP1 promoter region in 1205Lu-R treated with corin (2.5 μM, 24 h). **f,** Proliferation of 1205Lu-R cells overexpressing DUSP1 (DUSP1) treated with PLX4032 (72 h). Morphology of 1205Lu-R cells overexpression DUSP1 (bottom) compared to vector control (Ctrl), (top). Representative images shown, scale bar = 100 μm. Data (mean ± SE) are representative of at least two independent experiments (***p < 0.001, **p <0.01, *p <0.05).

As DUSP1 appeared to be a transcriptional target of CoREST, we next sought to determine whether DUSP1 transcriptional control regions were associated with the CoREST complex. Binding of the CoREST complex components HDAC1 and LSD1 to the promoter region of DUSP1 was analyzed by ChIP-qPCR, which confirmed the occupancy of both HDAC1 and LSD1 on DUSP1 promoter regions (Extended Data Fig. 9h). We next evaluated DUSP1 promoter accessibility in human melanoma cells following treatment with corin using ChIP-PCR analysis. H3K9ac and H3K4me2 active histone marks were found to be increased on the DUSP1 promoter following corin treatment, consistent with increased DUSP1 mRNA expression in response to corin (Fig. 3e).

To further explore the functional significance of DUSP1 expression in BRAFi-resistant melanoma, DUSP1 was constitutively expressed in BRAFi-R melanoma cells; conversely, DUSP1 expression was knocked down in BRAFi-sensitive melanoma cells using shRNA (Fig 3f, Extended Data Fig. 9i). 1205Lu-R cells overexpressing DUSP1 showed significant growth inhibition following PLX4032 treatment compared to vector control, suggesting that DUSP1 expression can resensitize BRAFi-R melanoma cells to PLX4032 (Fig. 3f). In addition, DUSP1 overexpression in 1205Lu-R cells resulted in distinct morphological changes resembling differentiated human melanocytes (Fig. 3f). In contrast, DUSP1 knockdown in human melanoma cells led to loss of PLX4032 sensitivity in BRAFi-sensitive melanoma cells (Extended Data Fig. 9j), further supporting a critical role for DUSP1 in mediating melanoma resistance to BRAFi treatment.

### Corin treatment of BRAFi-resistant melanoma promotes sensitization to PLX4032 in vivo

We next evaluated the effect of combining corin and PLX4032 in a BRAFi-R (1205Lu-R) melanoma mouse xenograft model. Mice with BRAFi-R tumors treated with PLX4032 alone showed progressive tumor growth that was similar to that of vehicle-treated animals (Fig. 4a). In contrast, corin alone significantly inhibited tumor growth in treated mice, while the combination treatment of corin + PLX4032 showed significantly greater inhibition of tumor growth compared to corin treatment alone (Figs. 4a-c). Notable necrotic areas (Fig. 4d) and decreased tumor cell proliferation (Fig. 4e) were observed in tumor specimens following corin treatment alone or in combination with PLX4032. Additionally, corin treatment with and without PLX4032 led to increased expression of H3K4Me2, H3K27Ac, DUSP1, and DUSP5, (Fig. 4f-i), as well as increased expression of markers of apoptosis and tumor hypoxia (Extended Data Fig. 10) in BRAFi-R tumors with associated decreases in expression of both MITF and AXL (Fig. 4j, k).

**Figure 4.**
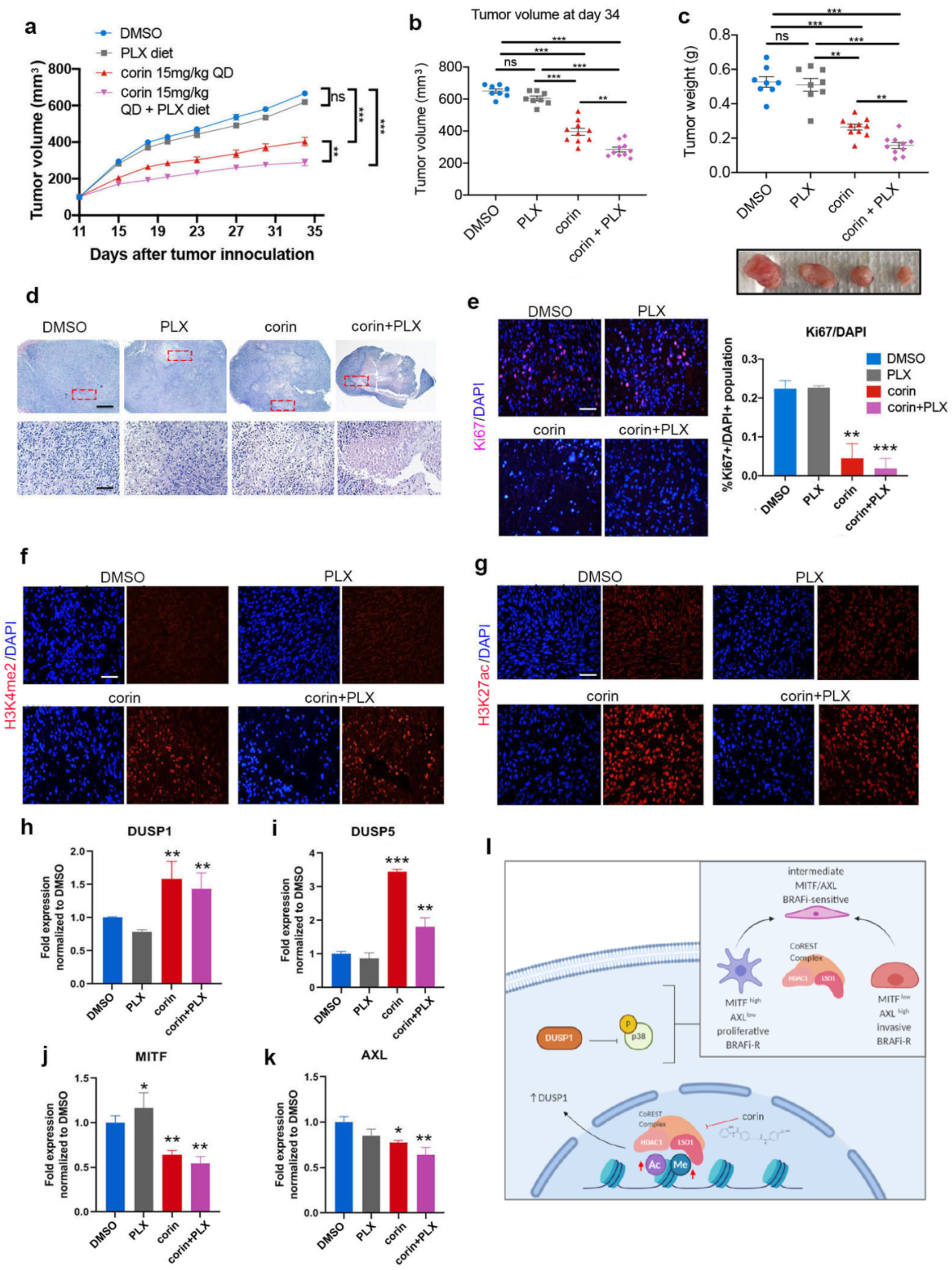
Corin treatment of BRAFi-R melanoma inhibits tumor growth and resensitizes tumors to PLX4032 in a mouse xenograft model. **a,** 1205Lu-R tumor growth in a mouse xenograft model following administration of corin (15mg/kg) +/− PLX4032. **b, c,** Tumor volume (**b**) and tumor weight (**c**) following treatment of 1205Lu-R xenografts with corin (15mg/kg) +/− PLX4032. Photo of representative tumors is included below. **d,**H&E staining of tumor xenografts showing areas of significant necrosis (light purple stain) in corin and corin+PLX treatment groups. (representative images shown, scale bar=1 mm (upper panel), 0.1 mm (lower panel)). **e,** Cell proliferation in corin +/− PLX4032 treated xenografts depicted by Ki67 staining and graphic quantification (representative images shown, scale bar = 100 μm). **f, g,** Immunofluorescence staining of H3K4me2 (**f**) and H3K27ac (**g**) expression in 1205Lu-R tumor xenografts following corin +/− PLX4032 treatment (representative images shown, scale bar = 100 μm). **h-k,** Expression of DUSP1 (**h**), DUSP5 (**i**), MITF (**j**) and AXL (**k**) in 1205Lu-R mouse xenografts treated with corin +/− PLX4032. Data (mean ± SE) are representative of at least two independent experiments (***p < 0.001, **p <0.01, *p<0.05). **l,** Model depicting the CoREST repressor complex as a central regulator of melanoma phenotype plasticity and resistance to targeted BRAFi therapies in human melanoma cells through repression of DUSP1.

## Discussion

Our results indicate that the CoREST repressor complex plays a critical role in promoting cellular plasticity and phenotype switching in melanoma as well as resistance to targeted BRAFi therapies through repression of the dual-specificity phosphatase, DUSP1. Furthermore, targeting of the CoREST complex with the small molecule inhibitor, corin, leads to increased expression of DUSP1, inhibition of p38 MAPK activity, and resensitization of BRAF-resistant melanomas to BRAFi therapy. Although global changes in the chromatin landscape have been noted during tumor progression in melanoma^8,9^, and histone H3 demethylases, including LSD1, have been shown to promote multidrug resistance^11,27^ and bypass of oncogene-induced senescence^28^, the specific targeting of these pathways has not led to meaningful clinical impacts to date. Here, we show that corin treatment of melanoma cells facilitates conversion of distinct melanoma phenotypes to an intermediate cell state (Fig. 1a-e) and that BRAFi-resistant melanomas show loss of distinctive cellular phenotypes and resensitization to PLX4032 following corin treatment (Figs. 1f-h, 2f, 4l).

Recent studies have defined the cellular plasticity and dynamic phenotypes associated with acquired MAPKi drug resistance in melanoma and epigenetic reprogramming observed in conditions of drug-induced stress that are ultimately fixed upon prolonged drug exposure and suggest that intermediate cellular phenotypes are notably drug-sensitive^4,29^. Here, we show that the CoREST repressor complex modulates these predominant MAPKi-resistant melanoma phenotypes including the differentiated/proliferative (MITF^high^/AXL^low^) and neural crest stem cell-like/invasive states (MITF^low^/AXL^high^) (Figs 1, 2), and that corin treatment of BRAFi-R melanomas promotes the emergence of an intermediate, BRAFi-sensitive phenotype. Such phenotype plasticity and associated drug-resistance has been widely studied during the process of epithelial-to-mesenchymal transition (EMT) in epithelial cancers, a process that is largely dictated by the expression of embryonic transcriptional programming^10^; however, melanomas possess a neural crest origin, suggesting that phenotype switch control mechanisms are likely to differ in this cell lineage^30^. As the REST/CoREST complex specifically regulates repression of differentiation-associated genes during neural development^14^, it is not wholly surprising that this complex would be reactivated to dictate the differentiation/proliferation balance in human melanoma cells. Indeed, expression of REST, the binding partner for CoREST, has been shown to be critical for early neural crest specification of developing melanoblasts^31^, a phenotype which resembles the undifferentiated/invasive/MITF^low^/AXL^high^ neural crest stem cell phenotype^29^. It is notable that RNA-seq data from two phenotypically distinct melanoma cell lines treated with the CoREST inhibitor, corin, showed lineage-specific effects on the expression of the EMT transcription factors ZEB1 and ZEB2, with downregulation of ZEB2 in the MITF^high^/AXL^low^/differentiated melanoma cells (451Lu-R) and downregulation of ZEB1 in the MITF^low^/AXL^high^/invasive/neural crest stem cells (1205Lu-R) consistent with corin suppression of the known distinct functions of these EMT-associated transcription factors in driving these specific melanoma phenotypes (Extended Fig. 2j)^32^. Upregulation of CDH1 in both melanoma phenotypes following corin treatment also suggests corin-associated inhibition of an EMT-like phenotype switch in these cells (Fig, 2d). Indeed, corin-associated phenotype changes in both MITF^high^ and AXL^high^ melanomas (Figs. 2a-e, Extended Data Figures 2-6) including increased expression of E-cadherin, and altered expression of MAPK, HIPPO, WNT signaling, and axonal guidance-associated genes are reminiscent of those seen following inhibition of the neural crest differentiation-associated SNAIL transcription factors during EMT in epithelial cancers. As the neural crest is an embryonic cell population of migratory and pluripotent cells that differentiate into diverse cell types including melanocytes, it is not surprising that cancer cells highjack these developmental programs for tumor initiation and progression.

Given the significance of SNAIL-regulated transcriptional programs during development, it would be expected that a higher order transcriptional network involving epigenetic reprogramming regulates the neural crest differentiation/migratory phenotype during development and the differentiation/proliferation phenotype in melanoma cells in a manner similar to EMT seen in epithelial cancers. Notably, the SNAG domain of SNAIL transcriptional repressor proteins has been shown to functionally associate with the CoREST repressor complex through the LSD1 domain, and this interaction is responsible for regulating EMT in breast cancers and hematologic malignancies^33^. Moreover, LSD1 inhibitors have been shown to disrupt LSD1/SNAG-domain protein-protein interactions which would similarly be expected to occur with the related analog, corin^34^. SNAIL transcriptional repressors have also been shown to positively regulate the expression of ZEB transcription factors^35^ which also associate with the CoREST complex during development^36^. We therefore hypothesize that CoREST mediates transcriptional programs governing melanoma phenotypes and BRAFi-resistance through interactions with EMT-associated transcription factors, including SNAIL and ZEB repressor proteins which are downstream of p38 MAPK^37^. Furthermore, we anticipate that such epigenetic hierarchical regulation is an essential feature of critical EMT-like embryonic and wound healing transcriptional programs as well as cellular responses to environmental stressors as seen in phenotype reprogramming of cancer cells exposed to pharmacologic therapies^38^. Indeed, a survey of melanoma cell lines using the Broad Institute Depmap portal (https://depmap.org/portal/) found statistically significant inverse dependencies of expression of MITF with SNAI1 and AXL with SNAI2 as well as a significant inverse relationship between SNAI1 and SNAI2 as well as ZEB1 and ZEB2. The Depmap survey also found significant direct relationships between SNAI1 and ZEB1 as well as SNAI2 and ZEB2 in melanoma cell lines, further supporting a potential role for SNAIL and ZEB transcription factors in the CoREST-mediated phenotype switch. Interestingly, our previous data demonstrated strong growth inhibition by corin in all melanoma and leukemia cell lines tested^5^; however, 50% of breast cancer cell lines and 70% of colon cancer cell lines evaluated also demonstrated significant growth inhibition by corin, suggesting that specific transcriptional programs, possibly relevant to a neural differentiation phenotype, may promote particular dependencies on CoREST-mediated cell growth.

As in development, chromatin structure in tumor cells is critical to cellular phenotype and cell fate. This process is quite malleable, as such plasticity lends itself to rapid reorganization under conditions of stress and serves as a cellular survival mechanism. Additionally, the rapid transitions afforded by epigenetic controls of gene expression serve as important mechanisms to evade therapeutic interventions^39^. Remarkably, we find that inhibition of CoREST activity in human melanoma specifically inhibits both distinctive proliferative and invasive cellular phenotypes and that such inhibition promotes resensitization of treatment-resistant melanoma cells to targeted BRAF inhibition. We further note that corin treatment of tumors in immunocompetent mice promoted antitumor immunity and impaired Foxp3+ Treg function^16^, suggesting that a therapeutic strategy targeting CoREST might enhance anticancer effects both through direct effects on tumor phenotype plasticity as well as an enhanced immune response (Extended Data Fig. 4). We therefore suggest that targeting of the CoREST complex in melanoma may provide a successful therapeutic approach to inhibit cellular plasticity and therapeutic resistance in human melanomas and other cancers that undergo similar phenotype transitions governed by epigenetic reprogramming.

**Supplementary Information** is included in a separate attachment.

## METHODS

### Cell Culture

Melanoma cell lines 451Lu, SKMel5, WM35, WM983B, Sbcl2, WM1552C, 1205Lu and A375 were obtained from Dr. Meenard Herlyn (The Wistar Institute, Philadelphia, PA). SKMel5 cells were obtained from Dr. Levi A. Garraway (Dana-Farber Cancer Institute, Boston, MA). SKMel5 BRAFi R cells were obtained from Dr. Deborah Lang (Boston University, Boston, MA). 451Lu BRAFi R and A375 BRAFi R cells were obtained from Dr. Jong-In Park (Medical College of Wisconsin, Wauwatosa, WI). All cell lines used were routinely checked for and found to be free of mycoplasma contamination. Melanoma cell lines were cultured in Dulbecco’s modified eagle medium (Invitrogen) supplemented with 10% fetal bovine serum, L-glutamine (2 mM), and 1% penicillin/streptomycin. All cell lines were maintained in a 37 °C incubator at 5% CO2.

### Compounds

PLX4032 (vemurafenib, #S1267) and MS275 (entinostat, #1053) were purchased from Selleck Chemicals. Corin and Cpd7 were provided by Dr. Philip Cole (Harvard Medical School, Boston, MA).

### Cell treatment with compounds

Corin, MS275, Cpd7 and PLX 4032 stocks were prepared in DMSO. Appropriate stock solutions were added to culture medium to achieve the desired final concentrations. An equal amount of DMSO was used as a vehicle control.

### Western blot

Whole-cell lysates were prepared in 3D-RIPA buffer. Proteins (20 μg) were separated by 10% or 12% SDS-PAGE and transferred to a polyvinylidene difluoride membrane. Membranes were blocked using 5% nonfat dry milk in PBS containing 0.05% Tween 20, and then incubated with primary antibody overnight at 4 °C. HRP-conjugated secondary antibody was used and detected using the Pierce ECL Western Blot Substrate. Antibodies were obtained from the following sources: H3K9ac (Abcam, ab32129, 1:1000), H3K4me2 (Abcam, ab32356, 1:5000), H3K27ac (Abcam, ab4729, 1:1000), Phospho-MEK1/2 (Cell Signaling, 9121, 1:1000), MEK1/2 (Cell Signaling, 9122, 1:1000), total H3 (Abcam, ab1791, 1:10000), DUSP1 (Millipore, 07-535, 1:1000), DUSP5 (Abcam, ab200708, 1:1000), Phospho-c-Jun (Ser63) (Cell Signaling, 9261s, 1:1000), c-Jun (Cell Signaling, 9165s, 1:1000), Phospho-Erk (Cell Signaling, 9101s, 1:1000), p44/42 MAPK (Erk1/2) (Cell Signaling, 9102s, 1:1000), MITF (Cell Signaling, 12590s, 1:1000), AXL (Cell Signaling, 8661s, 1:1000), Cleaved PARP (Cell Signaling, 9541s, 1:1000), Phospho-JNK (Cell Signaling, 4668s, 1:1000), JNK (Cell Signaling, 9252s, 1:1000), Phospho-p38 MAPK (Cell Signaling, 4631s, 1:1000), p38 MAPK (Cell Signaling, 9212s, 1:1000), GAPDH (Cell Signaling, 2118L, 1:2000), HRP-conjugated secondary antibodies (Santa Cruz Biotechnology, 1:5000 for β-actin blots, 1:2000 for all others). Blots shown are representative of at least two independent experiments.

### Western Blot Imaging and Densitometric Analysis

Chemiluminescent blots were imaged with the ChemiDoc MP imager (Bio-Rad). The Band Analysis tools of ImageLab software version 4.1 (Bio-Rad) were used to select and determine the background-subtracted density of the bands in all blots. The ratio of p-p38/p38 was calculated using densitometric readings of individual band of p-p38 and p38 for each treatment condition and normalized to control.

### PicoGreen^®^ cell proliferation assay

Cells were seeded in a 96-well plate and treated with inhibitor at the indicated concentrations. Media was removed and replaced with fresh media containing compound every 24 h. After 72 h incubation, 20 μl lysis buffer (10 mM Tris-HCl, 1 mM EDTA, 0.2% (v/v) Triton X-100) was added to each well and the plate was shaken vigorously on an orbit shaker for 10 min. 70 μl TE buffer was subsequently added to each well and the plate was agitated for 5 min. 95 μl of sample solution from each well was transferred to a new 96-well, non-clear bottom plate (NUNC 236105 96F) and 95 μl 1X PicoGreen^®^ solution (Life Technologies, P11496) was added to each well. The plate was then agitated for 5 min in the dark after which fluorescence was measured using excitation and emission wavelengths of 480 nm and 520 nm, respectively, using a SpectraMax microplate reader. Data represent at least two independent experiments with three technical replicates per experiment. Where *p*-values are reported, the unpaired *t* test was used to determine significance.

### CoREST1 and DUSP1 knockdown

shRNA clones targeting CoREST (TRCN0000128570 and TRCN0000129660) were obtained from the High Throughput Biology Center at Johns Hopkins University. DUSP1 MISSION shRNA Bacterial Glycerol Stock (SHCLNG-NM_004417, total five constructs were tested TRCN0000367616, TRCN0000367617, TRCN0000231489, TRCN0000002517, TRCN0000002516) was purchased from Sigma Aldrich.

Lentiviruses were produced in HEK293T cells using Lipofectamine^®^ 2000 (Invitrogen) according to the manufacturer’s instructions and stored at −80 °C after 0.22 μm filtration. For lentiviral infection, 451Lu BRAFi-R and 1205Lu BRAFi-R cells were incubated with CoREST shRNA. 451Lu BRAFi-sensitive and 1205Lu BRAFi-sensitive cells were incubated with DUSP1 shRNA or scramble containing lentiviral particles overnight. Cells were selected with puromycin 48 h after transduction to create stable cell lines. CoREST knockdown was determined by quantitative RT– PCR and confirmed by Western blot. shRNA sequences are provided in Supplementary Table 1.

### DUSP1 overexpression

DUSP1 human overexpression plasmid (Origene, NM_004417) was expanded and transfected into 451Lu BRAFi-R and 1205Lu BRAFi-R cells using jetPRIME® transfection reagent (Polyplus transfection). Cells were selected with kanamycin for 48 hours before drug treatment. DUSP1 transcript levels were determined by quantitative RT–PCR to confirm overexpression efficiency.

### Colony formation assay

451Lu BRAFi-R and 1205Lu BRAFi-R cells were seeded in 6-well tissue culture plates at 5000 cells per well and treated with DMSO as control, 2.5 μM of corin +/− 5μM PLX4032 or 5μM PLX4032 alone for 10 days. Following treatment, each well was fixed for 10 min with 4% paraformaldehyde (Electron Microscopy Services) diluted in PBS (Gibco) and then stained for 20 minutes with Crystal Violet (Fisher Chemical). After washing and drying, each well was photographed and quantified using ImageJ.

### Boyden Chamber Invasion Assay

Each 24-well 8.0 μm pore polycarbonate membrane Transwell® insert was coated with 50 μg Matrigel diluted in 30 μL serum free DMEM and allowed to polymerize at 37°C for 30 min (Corning). 15,000 cells pre-treated with DMSO, 1 μM corin, or 2.5 μM corin for 24 h were seeded in 300 μL serum-free media in the top chamber; 600 μL of 20% FBS DMEM was added to the bottom chamber. After 24 h of invasion, cells remaining in the top chamber were removed with a cotton swab and cells that invaded to the bottom of the membrane were fixed in 70% ethanol, stained with 50 μg/mL propidium iodide, and washed in PBS. Cut out membranes were mounted onto a slide with UltraCruz Mounting Medium (Santa Cruz Biotechnology). Four 10x and 20x images were acquired per membrane using the Nikon Eclipse E400 microscope and SPOT Advanced software. Fiji-Image J was used to count invaded cells using the protocol previously described^40^.

### Focal adhesion staining and quantification

Glass coverslips were placed into 6-well plates and 70,000 cells were plated in each well. Cells were treated with DMSO control or 2.5 uM corin treatment for 24 h. Next, media was aspirated and cells were fixed to slides with 4% paraformaldehyde in PBS for 10 min. Slides were then washed twice with 0.1% 100X triton in PBS. Blocking solution (10% goat serum in PBS with 0.25% 100X triton) was added for 30 min followed by anti-vinculin antibody (Millipore, MAB3574) diluted in blocking solution at 1:300. Slides were left to incubate on a rocker for 2 h at room temperature, and then washed twice with 0.1% 100X triton in PBS. Texas Red-X Phalloidin diluted 1:200 in PBS was added to each slide and left to incubate for 20 min at room temperature. Slides were washed twice with 0.1% 100X triton in PBS and mounted with Vectashield medium with DAPI (H-1800; Vector Labs). Quantification of focal adhesions was performed according to the protocol described previously^41^.

### RNA Sequencing

451Lu BRAFi-R and 1205Lu BRAFi-R cells were treated for 24 h with 5 μM PLX4032, 2.5 μM corin, 5 μM PLX4032 + 2.5 μM corin, or DMSO as a control. RNA was isolated from cells using an RNeasy Plus Mini kit (Qiagen Inc.) following the manufacturer’s instructions.

### RNA-Seq data analysis

Libraries were sequenced on Illumina one High Output and one Midi Output NextSeq 500 Next Generation Sequencing (~530M Single Reads). Reads were aligned to the GRCh37/hg19 using STAR (version v2.4.1c). Transcriptome assemblies and differential expression ratios were performed using featureCount: subread 1.4.6-p2 and voom-limma: 3.26.9. Genes were selected as following in all conditions: log2 Fold Change ≤−1 or ≥1; and nominal P-value of P value ≤ 0.05. Box plots and Volcano plots were generated using Prism 6 software (GraphPad Software).

### Publicly available database

DUSP1 expression in normal skin, benign nevi, and malignant melanoma was obtained from publicly available microarray data^42^. Kaplan-Meier curves were generated by GEPIA (http://gepia.cancer-pku.cn) using the TCGA database (http://cancergenome.nih.gov/). Comparison analysis was performed using publicly available whole-exome sequence (WES) datasets of serial tumor biopsies (baseline and acquired resistant tumors) from patients with advanced melanoma treated with MAPK inhibitor (MAPKi) regimens^22^, by comparing the list of ‘Loss of Function (LOF)’ and ‘Gain of Function (GOF)’ genes from the dataset with corin-upregulated and down-regulated genes.

### Quantitative real time-PCR

RNA was isolated from melanoma cells following the manufacturer’s instructions and cleaned using the RNeasy Mini kit (Qiagen Inc.). 1 μg of RNA was reverse transcribed using SuperScript^®^ III First-Strand Synthesis System kit (Invitrogen). Real-time quantitative PCR was performed for 40 cycles of 15 seconds at 95 °C and 30 seconds at 60 °C, using the Step One Plus Real-time PCR system (Applied Biosystems). Data represents two independent experiments with three technical replicates per experiment. The data was quantified using the delta (delta Ct) method. Where *p*-values are reported, the unpaired *t* test was used to determine significance. Transcripts were amplified using the primers listed in Supplementary Table 1.

### Chromatin immunoprecipitation assay (ChIP-PCR)

Melanoma cells (10 × 10^6^) were crosslinked in 1% formaldehyde for 7 min and lysed in cell lysis buffer (5 mM HEPES, 85 mM KCl, 0.5% NP-40, 1X protease inhibitor cocktail (Sigma), 1X phosphatase inhibitor cocktail (Sigma)) on ice for 20 min as previously described^5^. Cell nuclei were lysed in buffer containing 50 mM Tris-HCl, pH 8, 10 mM EDTA, 1% SDS and chromatin was fragmented using a Bioruptor (Diagenode) sonicator to obtain an average size of 250–300 base pairs. Chromatin concentration was measured by absorbance at 260nm in a Nanodrop spectrophotometer (1 unit/ml of chromatin corresponds to OD_260_ = 1) and diluted 1:10 with dilution buffer (165 mM NaCl, 0.01% SDS, 1.1% Triton X-100, 1.2 mM EDTA, 16.7 mM Tris-HCl, pH 8.0). 1 unit of chromatin was immunoprecipitated by protein A magnetic beads (Dynabeads, Invitrogen) after incubation with antibodies specific for HDAC1 (Abcam, ab19845), LSD1 (Abcam, ab129195), H3K9ac (Abcam, ab32129) and H3K4me2 (Abcam, ab32356). Normal goat IgG and non-antibody treated samples were used as negative controls. Following overnight immunoprecipitation, beads were washed twice consecutively with each of the following buffers: Lio-B (50 mM HEPES, pH 8.0, 140 mM NaCl, 1% Triton X-100, 0.1% sodium deoxycholate, 1 mM EDTA), Hio-B (50 mM HEPES, pH 8.0, 500 mM NaCl, 1% Triton X-100, 0.1% sodium deoxycholate, 1 mM EDTA), LiCl (10 mM Tris-HCl, 250 mM LiCl, 0.5% NP−40, 0.5% sodium deoxycholate, 1 mM EDTA) and TE (10 mM Tris-HCl, pH 8.0, 1 mM EDTA). Immunoprecipitated chromatin and input DNA were reverse-crosslinked in elution buffer (50 mM Tris-HCl, 10 mM EDTA, 1% SDS) in the presence of proteinase K (50 μg ml^−1^) by shaking (1300 RPM) at 68 °C for 5 h. DNA was purified using phenol-chloroform and precipitated in ethanol at −20 °C. DNA pellets were dissolved in 200 μL of ddH_2_O. The relevant primers are listed in Supplementary Table 1.

### Mouse melanoma xenograft

Mice were maintained under pathogen-free conditions in an American Association for Accreditation of Laboratory Animal Care (AAALAC)-accredited facility at the Boston University Medical Center, under the supervision of the Laboratory Animal Science Center (LASC) and its staff of veterinarians and support personnel. All animal studies were conducted in accordance with Boston University Institutional Animal Care and Use Committee Guidance.

For the 1205Lu BRAFi-R xenograft studies, six to eight-week old female Balb/C nude mice were purchased from Charles River Laboratories, Inc. (Wilmington, MA) and allowed to acclimate for 1 week prior to beginning the experiment. For each animal, 1205Lu BRAFi-R (6 × 10^6^) cells in 100 μl growth media mixed with 50% matri-gel (BD, USA) were injected bilaterally into the subcutaneous tissue of both flanks. On day 3 after tumor inoculation, animals were randomly assigned to two groups, which were administered vemurafenib diet or control diet by the Research Randomizer at http://www.randomizer.org. Vemurafenib diet (5.67 g/kg body weight to achieve a 100 mg/kg body weight daily dose) and control diet were prepared at Envigo (Madison, WI). When xenograft size reached an approximate volume of 100 mm^3^, the 20 mice were randomized into 5 mice per group with the average tumor volume distributed equally between groups. Vehicle control (5% DMSO/H_2_O) and corin (15 mg kg^−1^) were administered at 10 ml kg^−1^ IP once a day. The mice were maintained in a pathogen-free environment with free access to food and water. Body weight and tumor volume were measured twice weekly. Tumor size was measured with linear calipers and calculated using the formula: ([length in millimeters × (width in millimeters)^2^]/2). The mice were sacrificed after 23 days and tumor weights were measured. Three animals from each treatment group were chosen at random for follow-up data analysis. Where *p*-values are reported, the unpaired *t* test was used to determine significance.

### Xenograft tumor sample processing and immunohistochemistry

After sacrificing mice, tumors were removed and fixed in 3.7% formaldehyde. Serial sections 5 μm thick were cut from the formalin fixed, paraffin embedded tissue blocks, floated onto charged glass slides (Super-Frost Plus, Fisher Scientific), and dried overnight at 60 °C.

Sections were deparaffinized and hydrated using graded concentrations of ethanol to deionized water prior to immunohistochemistry. Tumor sections were blocked in serum (5% serum in PBS-T (0.5% TritonX-100 in PBS)), and then incubated overnight at 4°C in Ki 67 (Abcam, ab15580, 1:500), Cleaved Caspase-3 (Cell Signaling, 9661s, 1:500), HIF1 alpha (Abcam, ab16066, 1:500). Sections were then incubated with fluorescence-conjugated (FITC) goat anti-rabbit secondary antibody for 1 h at room temperature, washed with PBS, and mounted using VECTASHIELD mounting medium (Vector Laboratories).

### H&E staining

Animal tissue H&E staining was performed at Applied Pathology in Worcester, MA using their standard protocol (https://files.sitebuilder.name.tools/enom52979/file/hestainingprotocol.pdf).

### Statistical significance

Statistical significance was tested either using 2-tailed Student *t*-test or the Wilcoxon rank–sum test followed by a Benjamini–Hochberg correction for false discovery rate. Results were considered significant for adjusted *P* < 0.05.

### Calculation of the combination index (CI)

The combined activity of corin and PLX4032 was determined by calculating the CI for both compounds in 451Lu-R and A375-R cells in Compusyn software^18^. The concentrations used for PLX4032 alone are: 0.01μM, 0.1μM, 1μM, 5μM, 10μM; concentrations used for corin alone are: 0.01μM, 0.1μM, 1μM, 5μM, 10μM. Concentrations used for combination treatment are: 1μM PLX4032 + corin (0.01μM, 0.1μM, 1μM, 5μM, 10μM). The results are interpreted as: 0 < CI < 1 indicates synergism; CI = 1 indicates an addictive effect; and CI >1 indicates antagonism.

## Data Availability

RNASeq data have been deposited in the NCBI GEO database with accession number GSE156250. The data that support this study are available from corresponding authors upon request.

## Code availability

The code to reproduce the mRNA-Seq analysis and plots is publicly available at https://bitbucket.org/bucab/corin_melanoma

## Acknowledgements

M.W. is supported by a Karin Grunebaum Cancer Foundation Research Award. P.C. is supported by NIH R37GM62437. R.A. is supported by a Melanoma Research Alliance Established Investigator Award (MRA 622586).

## Author Contributions

M.W., P.C., and R.A. conceived the study, interpreted data and wrote the manuscript. M.W., A.H., F.G., K.K., S.N., M.C., M.C., and A.X. carried out the cell-based work. M.W. and K.K. carried out the mouse xenograft studies. J.K. designed and synthesized the CoREST inhibitor, corin. A.L., F.A., and M.W. performed RNA-seq data analysis. All authors were involved in the experimental design, data interpretation, and manuscript editing.

## Materials & Correspondence

Supplementary Information is available in the online version of the paper.

Correspondence and requests for materials should be addressed to M.W, P.A.C, R.M.A.

