## Supplemental Files/Extended Data for "The CoREST Repressor Complex Mediates Phenotype Switching and Therapy Resistance in Melanoma"

Table of Contents:


I. Extended Data Figures and Tables S2-S11

II. Supplementary Table S12

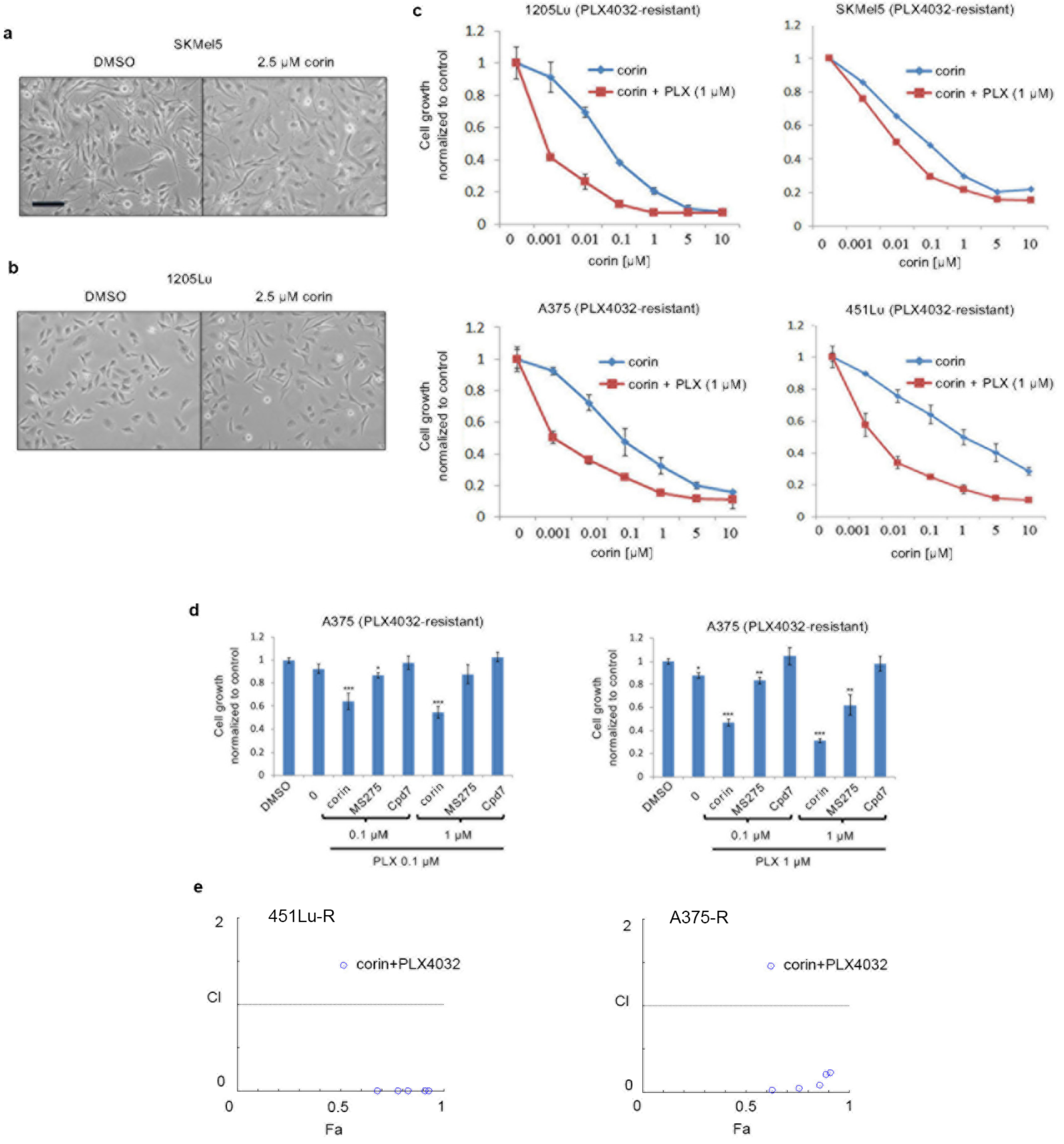


**Extended Data Figure 1 | CoREST inhibition of melanoma cells promotes changes in cell morphology and resensitization to PLX4032 in BRAF-R melanoma cells**. a, b, Morphological changes in SKMel5 (a) and 1205Lu cells (b) following treatment with 2.5 µM corin for 24 h. c, Proliferation assays of 1205Lu-R and A375-R ($MITF^{low}/AXL^{high}$) cell lines and SKMel5-R and 451Lu-R ($MITF^{high}/AXL^{low}$) cell lines treated with increasing doses of corin +/- 1 µM PLX4032 for 72 h. d, Proliferation assays of A375-R cells treated with PLX4032 0.1 µM (left panel) and 1 µM (right panel)) +/- corin, MS275 or Compound7 (Cpd7) (0.1 µM and 1 µM). e, Combination Index (CI) measurements for corin and PLX4032 show strong synergy for two BRAFi-R melanoma lines (451Lu-R and A375-R). Plots indicate the degree of synergy where >1 indicates antagonism, 1 indicates additivity, and <1 indicates synergy vs. the Fa=fraction affected (cell proliferation inhibition) across a range of PLX and corin concentrations. Representative images shown, scale bar = 100 µm. (***$p < 0.001$, **$p < 0.01$, *$p < 0.05$).

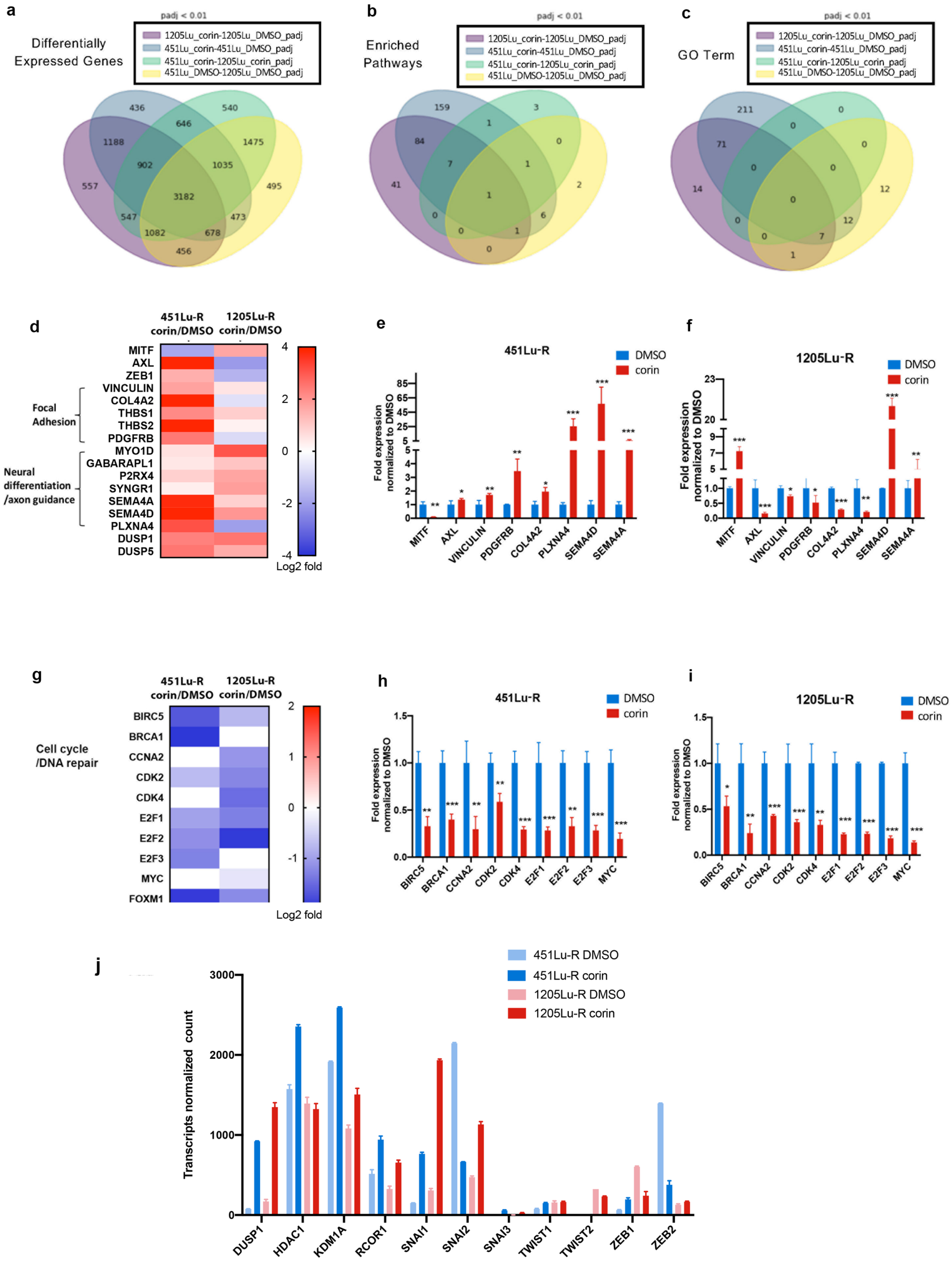


**Extended Data Figure 2 | Corin differentially regulates cell adhesion, neural differentiation, cell cycle, and DNA repair pathways in 451Lu-R vs 1205Lu-R melanoma cells.** a, b, c, Venn diagrams of RNA-seq data depicting a, differentially expressed genes (a), enriched pathways (b), and enriched gene ontology (GO) terms (c) that overlap between treatment groups. d, g, heatmaps illustrating gene expression patterns associated with focal adhesion and neural differentiation/axon guidance (d), and cell cycle/DNA repair (g) in 451Lu-R vs 1205Lu-R following corin treatment (2.5 µM, 24 h). e, f, and h, i, qPCR validated expression of differentially expressed genes associated with focal adhesion and neural differentiation/axon guidance and cell cycle/DNA repair, respectively, in 451Lu-R vs 1205Lu-R following corin treatment (2.5 µM, 24 h). j, Normalized transcript counts of EMT transcription factors in 451Lu-R vs 1205Lu-R following corin treatment (2.5 µM, 24 h). (***$p < 0.001$, **$p < 0.01$, *$p < 0.05$).

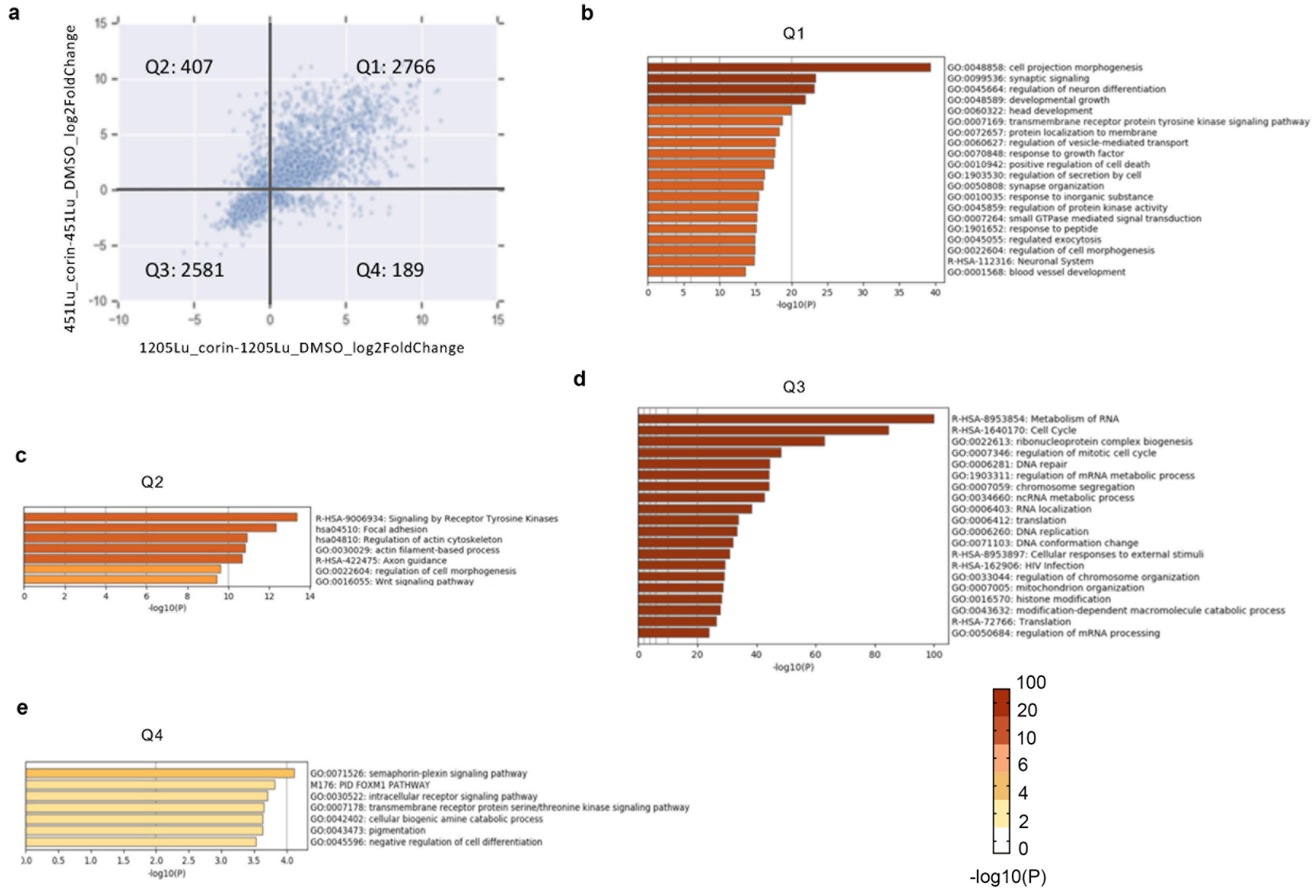


**Extended Data Figure 3 | GSEA illustrating GO classifications differentially expressed following corin treatment of 1205Lu-R and 451Lu-R cell lines.** a, Common DEGs (differentially expressed genes, fold change ≥ 2σ, FDR < 0.01) in 1205Lu-R and 451Lu-R cells treated with corin (2.5 µM, 24 h). were plotted into quadrants. Q1, up in both; Q2, up in 451Lu-R, down in 1205Lu-R; Q3, down in both; Q4, up in 1205Lu-R, down in 451Lu-R. GSEA associated with individual quadrants b, Q1; c, Q2; d, Q3;e, Q4.

**a**

451Lu-R corin vs DMSO
Reactome Interferon gamma signaling

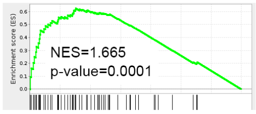


Top leading edge genes of Reactome Interferon gamma signaling in 451Lu-R

| | 451Lu-R corin vs. DMSO log2FoldChange | 451Lu R corin vs. DMSO_padj |
|---|---|---|
| HLA-DRA | 9.622 | 3.30E-10 |
| CAMK2B | 7.493 | 6.72E-06 |
| OASL | 7.044 | 4.06E-05 |
| HLA-DRB1 | 6.251 | 5.14E-08 |
| TRIM29 | 5.935 | 2.31E-03 |
| GBP5 | 4.834 | 9.49E-03 |
| HLA-DQA2 | 4.798 | 4.82E-02 |
| HLA-DRB5 | 4.778 | 5.05E-02 |
| HLA-DPB1 | 4.483 | 3.99E-05 |
| HLA-DQA1 | 4.116 | 4.64E-02 |

**b**

1205Lu-R corin vs DMSO
Reactome Interferon gamma signaling

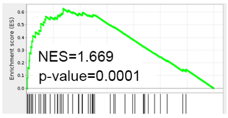


Top leading edge genes of Reactome Interferon gamma signaling in 1205Lu-R

| | 1205Lu-R corin vs. DMSO log2FoldChange | 1205Lu-R corin vs. DMSO_padj |
|---|---|---|
| HLA-DRB1 | 6.174933468 | 7.46E-04 |
| HLA-DQB1 | 6.051579228 | 2.23E-04 |
| GBP5 | 4.750462091 | 1.89E-04 |
| TRIM48 | 4.709467957 | 5.10E-02 |
| TRIM10 | 4.211435964 | 1.18E-01 |
| GBP2 | 4.018569222 | 1.05E-41 |
| HLA-DRA | 3.523307355 | 4.28E-17 |
| HLA-DPA1 | 3.349360326 | 3.66E-02 |
| HLA-DPB1 | 3.281761828 | 1.65E-01 |
| IRF5 | 3.135721973 | 1.06E-02 |

**c** GSEAs analysis of type I and type II IFN pathways

| | 451Lu-R corin vs. DMSO_pval | 451Lu-R corin vs. DMSO_NES | 1205Lu-R corin vs. DMSO_pval | 1205Lu-R corin vs. DMSO_NES |
|---|---|---|---|---|
| Reactome Interferon gamma signaling (type II) | 0.0001001 | 1.665346666 | 0.00010001 | 1.66943689 |
| Reactome Interferon alpha/beta signaling (type I) | 0.271467584 | 1.11842868 | 0.777811306 | 0.867429259 |

| | 451Lu-R PLX_corin vs. PLX_pval | 451Lu-R PLX_corin vs. PLX_NES | 1205Lu-R PLX_corin vs. PLX_pval | 1205Lu-R PLX_corin vs. PLX_NES |
|---|---|---|---|---|
| Reactome Interferon gamma signaling (type II) | 1.0036E-05 | 1.76312354 | 1.00033E-05 | 1.564305462 |
| Reactome Interferon alpha/beta signaling (type I) | 0.027238266 | 1.339255357 | 0.972307229 | 0.659322288 |

**d** Transcripts changes of RISC components and repetitive elements induced by corin

| | 451Lu-R corin vs. DMSO log2FoldChange | 451Lu-R corin vs. DMSO_pvalue | 1205Lu-R corin vs. DMSO log2FoldChange | 1205Lu-R corin vs. DMSO_pvalue |
|---|---|---|---|---|
| DICER1 | -0.254 | 5.38E-02 | -0.234 | 9.94E-02 |
| AGO2 | -0.674 | 2.59E-10 | -0.816 | 3.19E-30 |
| ERV1 | 0.336388006 | 1.27703E-47 | 0.751130744 | 0 |
| ERVL | 0.286339284 | 3.73609E-26 | 0.4376894 | 4.11635E-64 |

**Extended Data Figure 4 | Corin treatment of melanoma cells promotes expression of gene signatures associated with the type 1 and type 2 interferon response, and repetitive elements while decreasing expression of genes associated with the RNA-induced silencing complex (RISC)**. a, b, GSEA analysis in corin (2.5 µM, 24 h) versus DMSO-treated melanoma cells illustrating enriched Interferon gamma signaling in 451Lu-R (a) and 1205Lu-R (b) cells. NES (Normalized Enrichment Score), (FDR < 0.05). Gene expression changes in the top leading edge genes of the interferon gamma signaling pathway in 451Lu-R (a) and 1205Lu-R (b) cells. c, GSEA analysis of type I and type II Interferon signaling pathways in corin versus DMSO treated samples, and PLX4032 + corin versus PLX4032 alone treated samples. d, Expression of RISC component genes, and repetitive element genes in 451Lu-R and 1205Lu-R melanoma cells following treatment with corin .

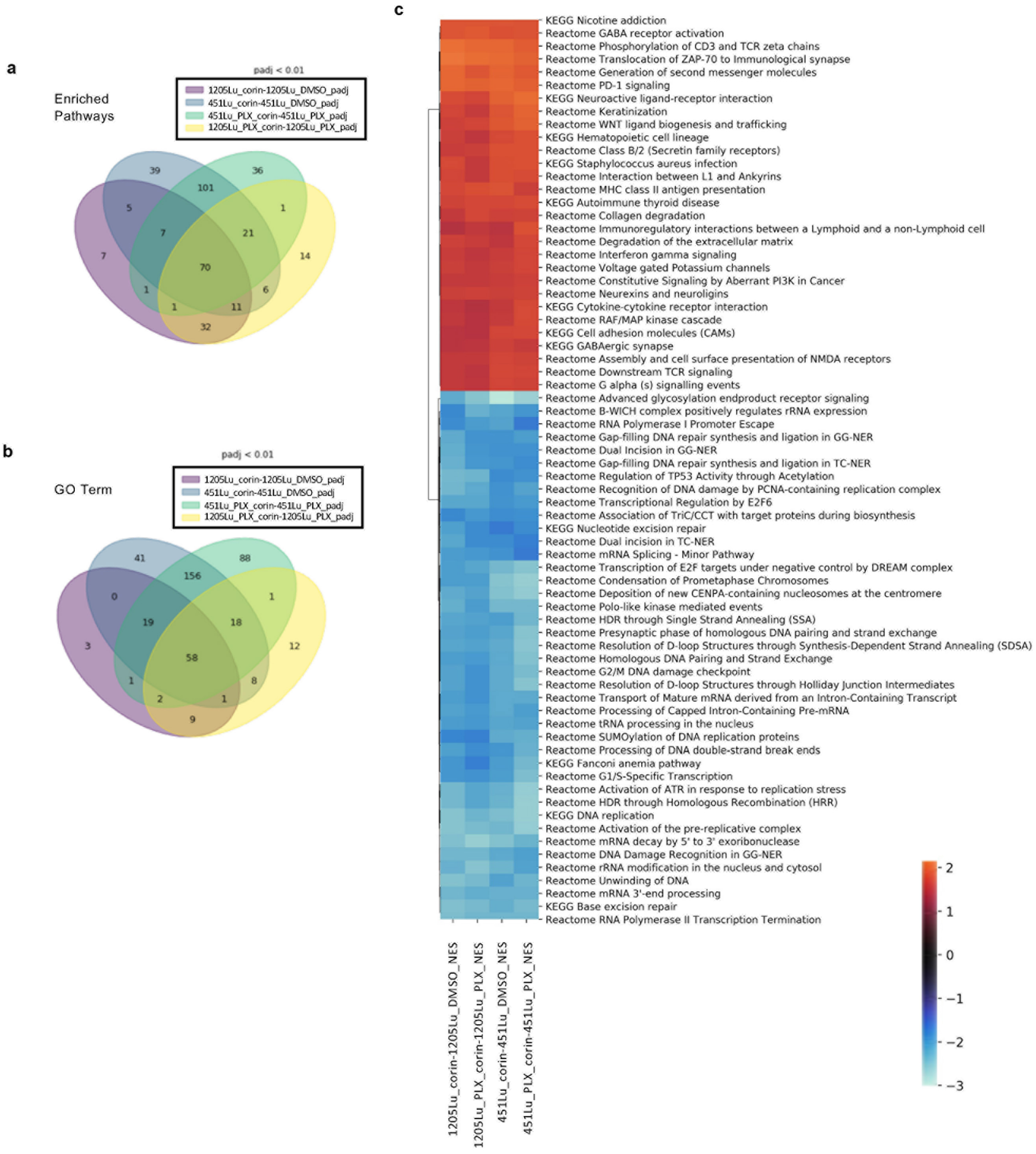


**Extended Data Figure 5 | Corin treatment of melanoma cells induces common KEGG and Gene Ontology pathway changes in 1205Lu-R and 451Lu-R cell lines**. Venn Diagram depicts a, Enriched KEGG pathways and b, enriched Gene Ontology (GO) terms that overlap between corin (+/- PLX4032) treatment groups. c, heatmap of GSEA common corin-induced pathways in 1205Lu-R and 451Lu-R melanoma cells.

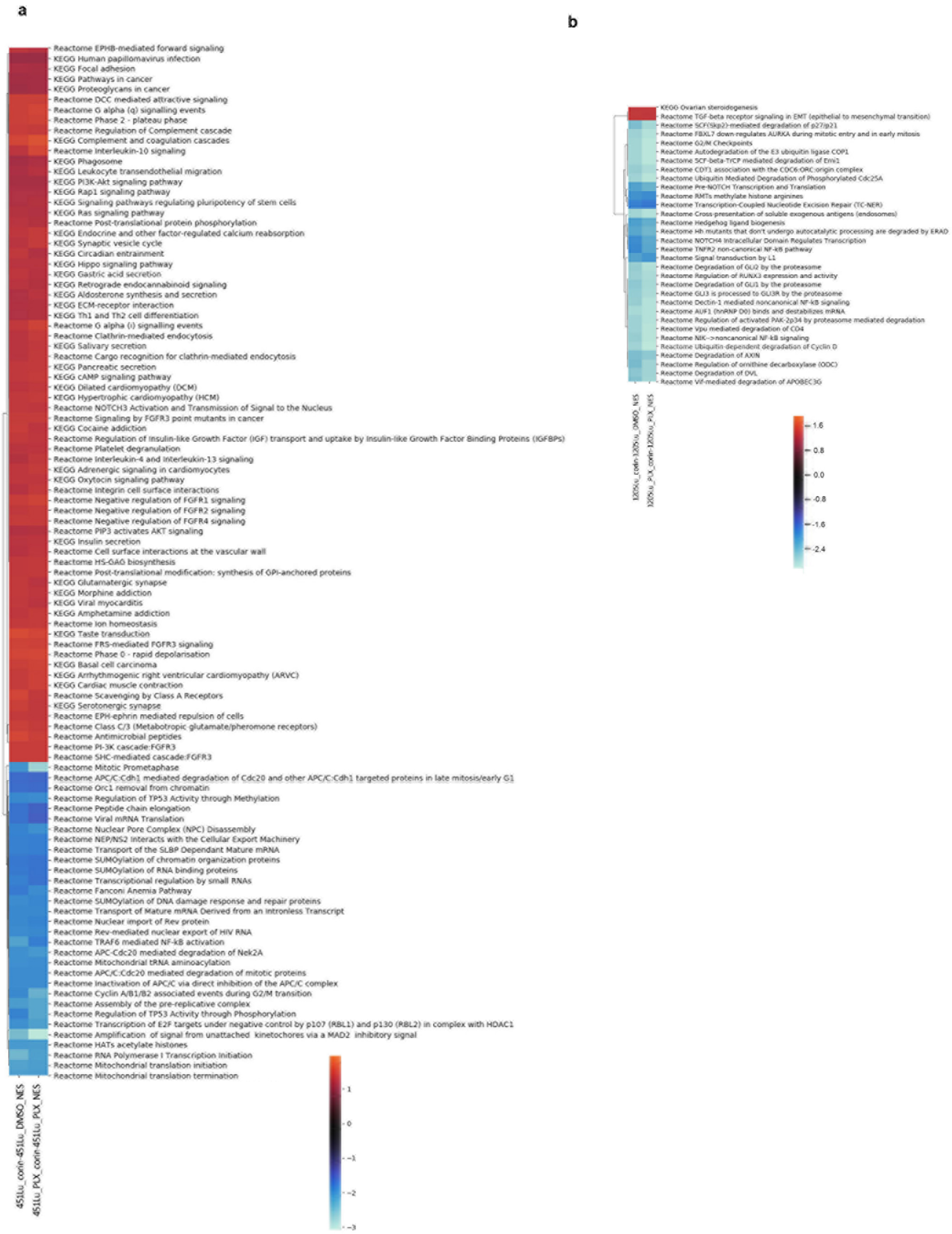


**Extended Data Figure 6 | Corin induces unique GSEA pathways in BRAFi-resistant cell lines.** Heatmaps of unique enriched pathways in a, 451Lu MITF$^{high}$/AXL$^{low}$ BRAFi-resistant cell lines and b, 1205Lu MITF$^{low}$/AXL$^{high}$ BRAFi-resistant cell lines treated with 2.5 μM corin +/- PLX4032 for 24 h.

a

**Expression of Genes Downregulated in Patient Melanomas During Acquired MAPKi Resistance**

| | 451Lu-R (PLX+corin)/PLX | 1205Lu-R (PLX+corin)/PLX |
|---|---|---|
| Gene | Log2fold change | |
| AXIN2 | 3.02 | 3.22 |
| CRLF2 | 7.52 | 4.13 |
| CTNNA2 | 6.53 | 3.87 |
| FGF12 | 5.93 | 2.29 |
| FGF5 | 7.11 | 6.46 |
| FGFR2 | 2.97 | 5.63 |
| FOXO1 | 2.63 | 2.90 |
| MMP1 | 13.73 | 4.02 |
| NR4A1 | 3.30 | 4.19 |
| PLD1 | 2.06 | 2.00 |
| RUNX1T1 | 6.42 | 2.82 |
| SIRT4 | 2.32 | 3.32 |
| WNT11 | 7.47 | 6.98 |

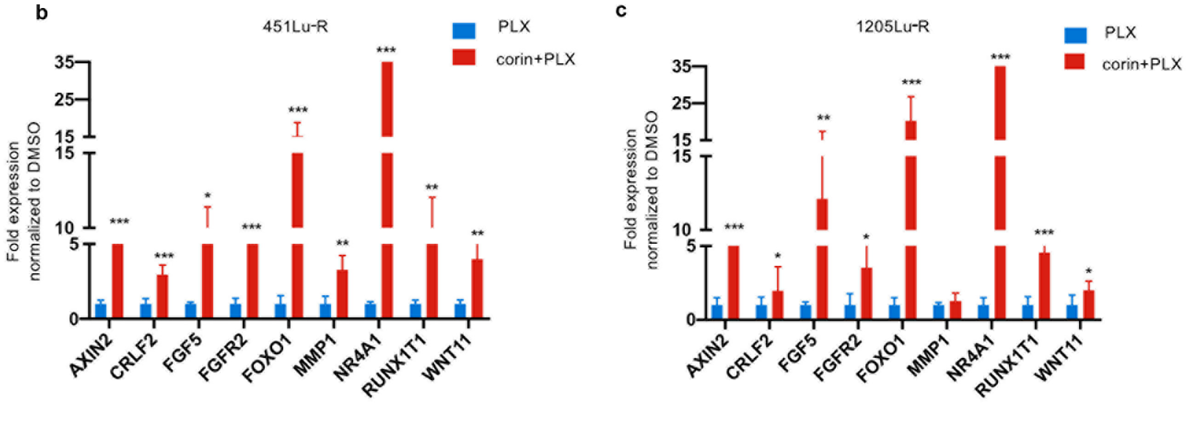


**Extended Data Figure 7 | Genes downregulated during the acquisition of MAPKi-resistance in melanoma patients[22] are induced following corin treatment.** a, Table of genes repressed in MAPKi-resistant melanoma specimens and associated level of induced expression following 24 h treatment with corin + PLX4032 in 451Lu-R and 1205Lu-R melanoma cells. b, c, qPCR validation of genes downregulated during acquired MAPKi-resistance in human melanoma specimens which are induced following 24 h treatment with corin + PLX4032 vs. PLX4032 alone in 451Lu-R (b) and 1205Lu-R (c) cells. (***p < 0.001, **p <0.01, *p<0.05).

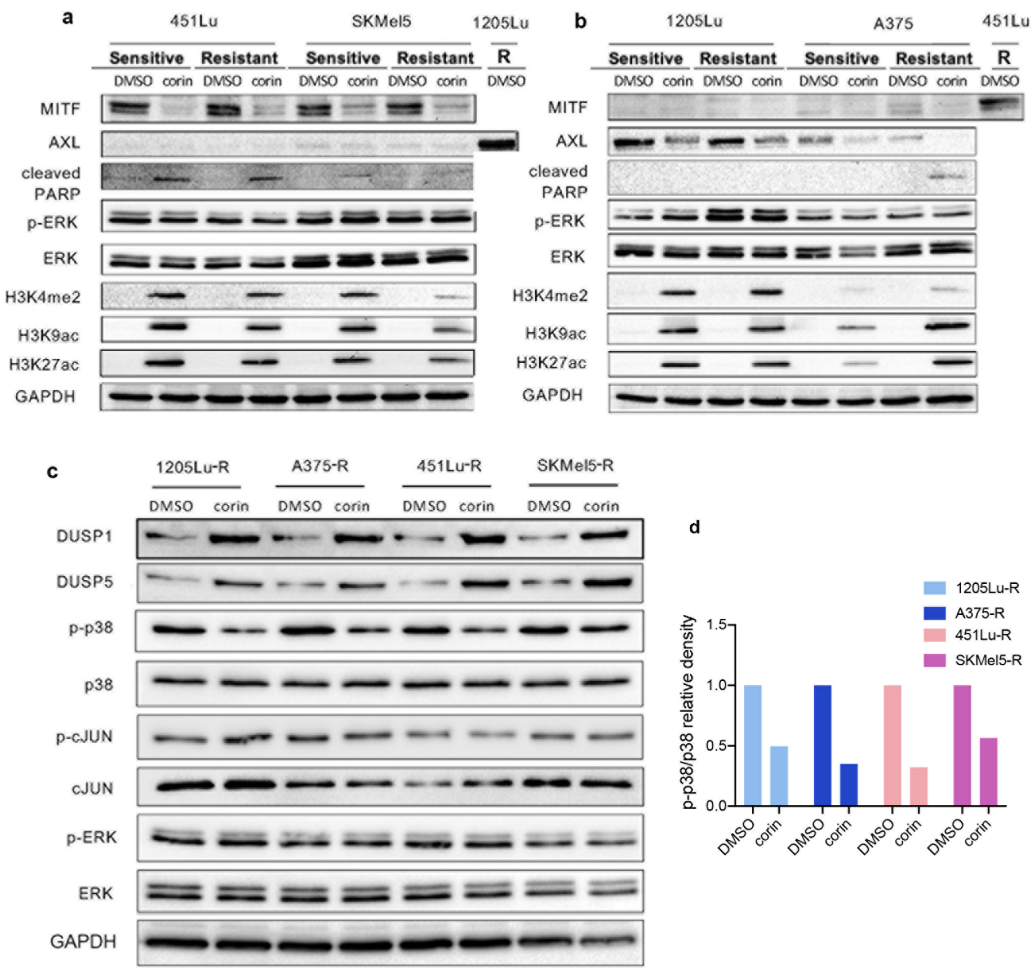


**Extended Data Figure 8 | Corin mediates phenotype switching and re-sensitization to BRAFi therapy through DUSP1-associated inhibition of p38 MAPK in melanoma cells**. a, b, Western blot analysis of MAPK pathway activity in BRAFi-sensitive and -resistant melanoma cell lines treated with 2.5 µM corin for 24 h (a), 451Lu and SKMel5 cell lines and (b), 1205Lu and A375 cell lines. c, Western blot analysis of DUSP expression and MAPK-associated proteins in 1205Lu-R, A375-R, 451Lu-R and SKMel5-R cells treated with 2.5 µM corin for 48 hours. d, Quantification of relative expression of pp38 (active) versus p38 (total) protein expression in 1205Lu-R, A375-R, 451Lu-R and SKMel5-R melanoma cells following treatment with corin.

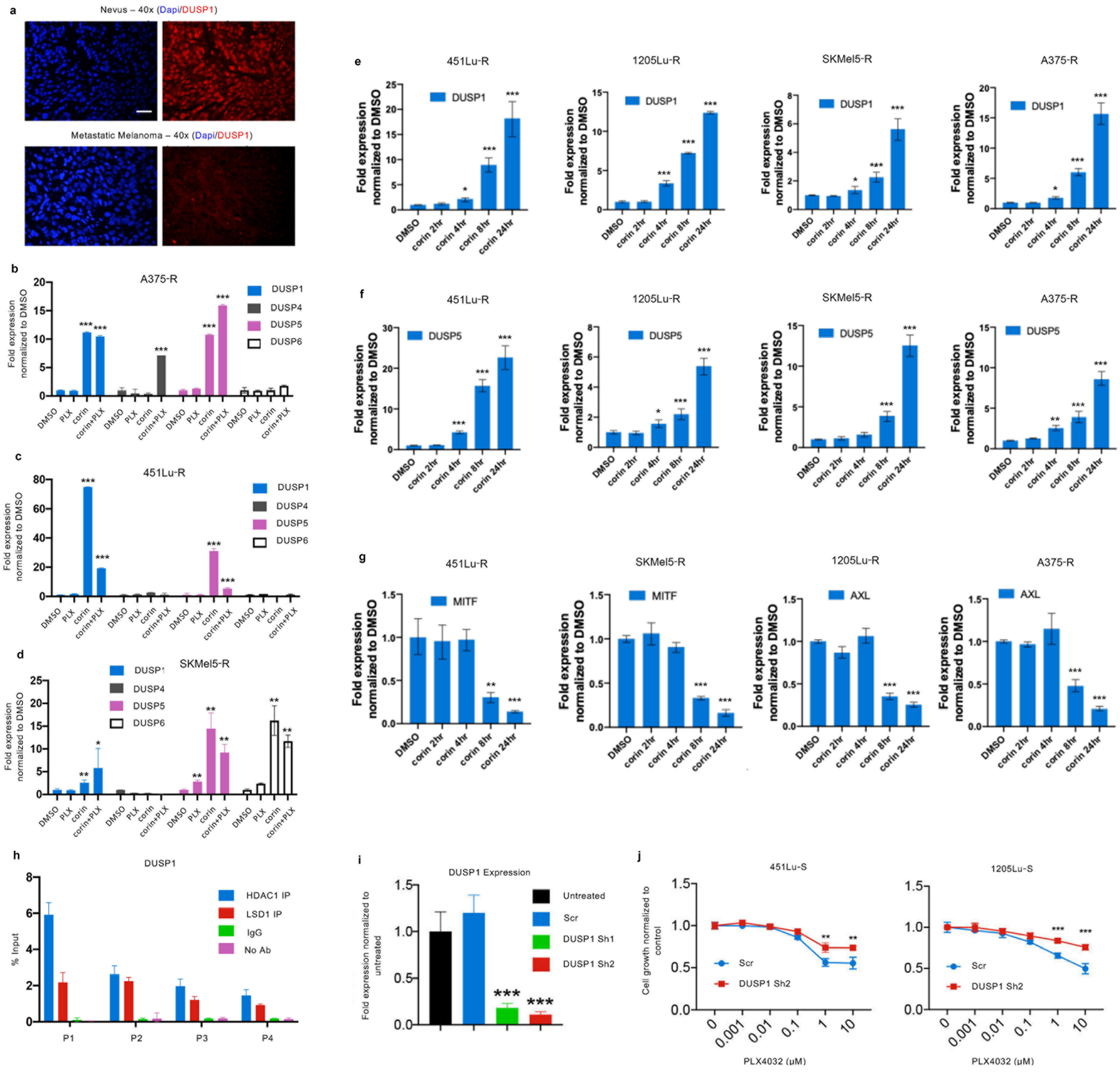


**Extended Data Figure 9 | Corin induces expression of DUSP1 in BRAFi-resistant melanoma cells which promotes senstivity to PLX4032 and is downregulated in human melanomas versus benign nevi.** a, Immunostain of DUSP1 expression in human melanoma vs. benign nevus. Representative images shown, scale bar = 100 µm. qPCR analysis of DUSP1, 4, 5, and 6 expression in b, A375-R, c, 451Lu-R, and d, SKMel5-R cells following 24 h treatment with 5 µM PLX4032, +/- 2.5 µM corin. e-g, qPCR analysis of DUSP1 (e), DUSP5 (f), MITF and AXL (g) in 451Lu-R, 1205Lu-R, A375-R, and SKMel5-R cells following treatment with 2.5 µM corin. h, Binding of the CoREST complex components (HDAC1 and LSD1) to the promoter regions of the DUSP1 gene by ChIP-qPCR. IgG and No Ab conditions were used as control. i, DUSP1 knockdown by shRNA lentivirus in 1205Lu cells, confirmed by qPCR. j, Cell growth assay of 451Lu and 1205Lu melanoma cells treated with PLX4032 (72 h) following knockdown of DUSP1 versus scramble control, (n=3). (***p < 0.001, **p <0.01, *p <0.05).

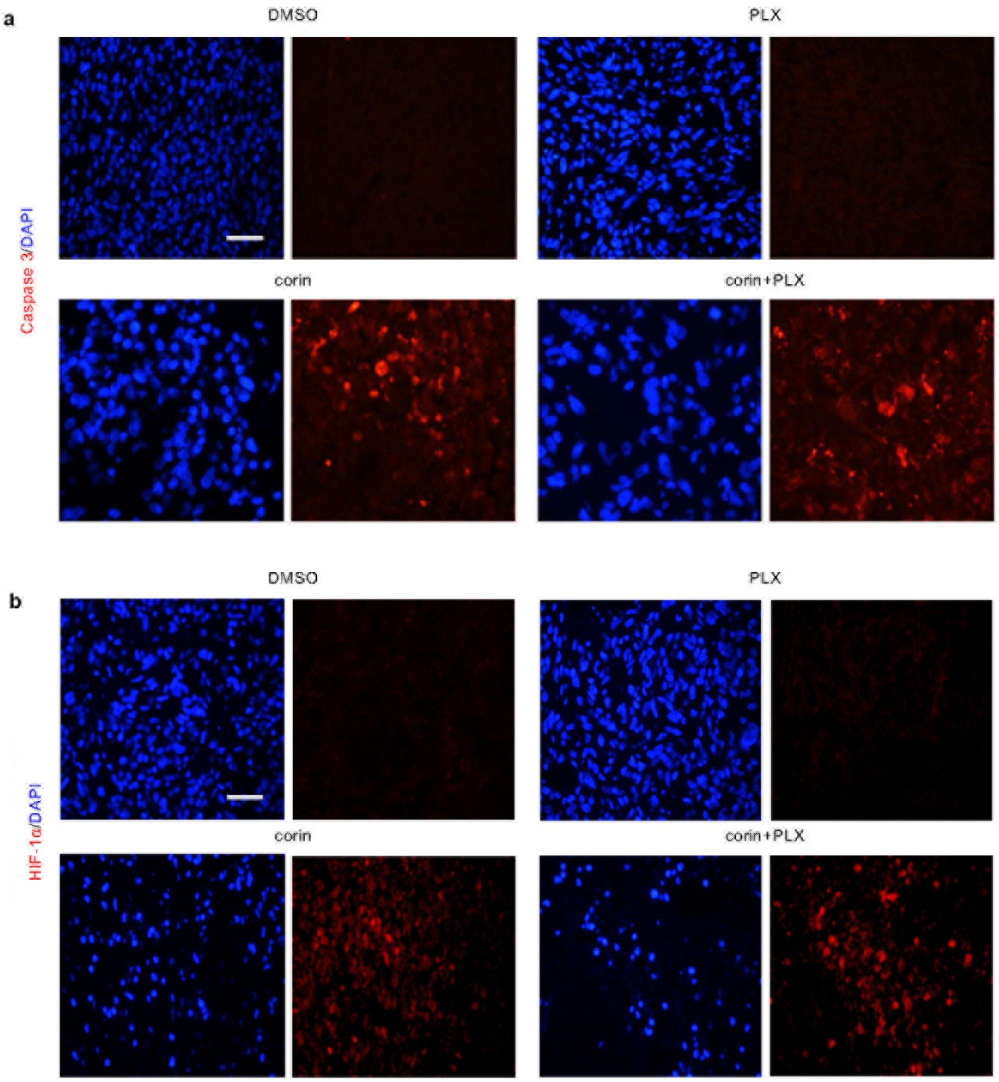


**Extended Data Figure 10 | Corin promotes apoptosis and hypoxia in BRAFi-resistant melanoma xenografts.** Immunofluorescence stain of a, cleaved caspase 3 and b, HIF-1α in 1205Lu-R melanoma xenografts treated with DMSO, PLX4032 alone, corin alone, corin + PLX4032. Representative images shown, scale bar = 100 μm.

**Supplementary Table 1 | Primer sequences used for RT-qPCR and ChIP-qPCR.**

| RT-qPCR Primer Sequences | | |
|---|---|---|
| **Target** | **Forward Primer (5'->3')** | **Reverse Primer (5'->3')** |
| AXIN2 | GGATCACTGGCTCCGCGA | AGTTCCTCTCAGCAATCGGC |
| AXL | CCAGGACACCCCAGAGGTGCTAAT | TGGTGGACTGGCTGTGCTTGC |
| BIRC5 | CCACTGAGAACGAGCCAGACTTG | AGAAAGGAAAGCGCAACCGG |
| BRCA1 | GAATTTATCGAGTGGCCAAAC | TCAAAGACTTGACTGTTGTGG |
| CCNA2 | GCATGTCACCGTTCCTCCTT | GGGCATCTTCACGCTCTATTT |
| CDK2 | GCTTTTGGAGTCCCTGTTCG | GGTCCCCAGAGTCCGAAAGA |
| COL4A2 | CCTGAAGGCACAGCTAACCA | TGCTGTTGTCTCGTCTGTCC |
| CRLF2 | ACTCCTGTTTCAGGCATGGG | AGTCAGGTTGGTCCTGGAGT |
| DUSP1 | AGGACAACCACAAGGCAGAC | CAGTGGACAAACACCCTTCC |
| DUSP4 | CAAAGGCGGCTATGAG | GGTTATCTTCCACTGGG |
| DUSP5 | GCCCGCGGGTCTACTTCCTC | CTCGGAGGTCCGTCGGGAGA |
| DUSP6 | CGAGACCCCAATAGTGC | AATGGCCTCAGGGAAA |
| E2F1 | AGGGGTGTGGGGTTGATACC | TCAGACACTGCAGGAGGGAC |
| E2F2 | CCTTGGAGGCTACTGACAGC | CCACAGGTAGTCGTCCTGGT |
| FGF5 | CTCTCTCTTCCCCTCTCCCC | TAGGGCTGATTCTGGGCTCT |
| FGFR2 | CCTGCGGAGACAGGTAACAG | GGTGTCTGCCGTTGAAGAGA |
| FOXO1 | GATCCCGTAAGTCGGGCGG | GCTGCTGCCTGTTGAATGTG |
| GAPDH | GGCTCTCCAGAACATCATCCCTGC | GGGTGTCGCTGTTGAAGTCAGAGG |
| MITF | GGAAATCTTGGGCTTGATGGA | CCCGAGACAGGCAACGTATT |
| MMP1 | AAGGCCAGTATGCACAGCTT | GGGCCACTATTTCTCCGCTT |
| MYC | TCACCAGCACAACTACGCCG | CAGGATGTAGGCGGTGGCTT |
| NR4A1 | GCTACGAAACTTGGGGGAGT | ATGTGGCTTGCACCTGTTCT |
| PDGFRB | CAAGGACACCATGCGGCTTC | AGCAGGTCAGAACGAAGGTG |
| PLXNA4 | AATTCCCGTCTTCACGGAGG | AGTCACTGTTGCTCTGTGGG |
| RUNX1T1 | GTCAAGAAGCAGACCGGGAA | CTAGTGCAACTGGGTCTGGG |
| SEMA4A | CGGGATGGGGTTGAGAATGG | ATCATGCCAAAGCCAGCACT |
| SEMA4D | CTGGGGCTCATCTCTAGCAC | CTCTCACCACCGCAATGTCA |
| VCL | GCTGCTGTTAATGCCATCCAA | CCAACCGAGTCACCTCATCT |
| WNT11 | GTGAAGGACTCGGAACTCGT | GTCGCTTCCGTTGGATGTCT |
| **ChIP-qPCR Primer Sequences** | | |
| **Target** | **Forward Primer (5'->3')** | **Reverse Primer (5'->3')** |
| DUSP1 P1 | TGACTGTCCTATCTTTCCATGTTC | AAACATCGTCCCTGGCAAC |
| DUSP1 P2 | AATCTGTTACCACCTCTCTTTTTATTT | GTGGAGTCAAAGGAGGCAAA |
| DUSP1 P3 | TCATGCCTGATATTAAAGGTAGTCC | CCTCACTAGCAGAGAACACACAA |
| DUSP1 P4 | CTCCCAAAGTGCTGGGATTA | TTCTGAAGACACTTCCAAAATAGAA |
